# Ankyrin-R regulates fast-spiking interneuron excitability through perineuronal nets and Kv3.1b K^+^ channels

**DOI:** 10.1101/2021.01.21.427626

**Authors:** Sharon R. Stevens, Colleen M. Longley, Yuki Ogawa, Lindsay H. Teliska, Anithachristy S. Arumanayagam, Supna Nair, Juan A. Oses-Prieto, Alma L. Burlingame, Matthew D. Cykowski, Mingshan Xue, Matthew N. Rasband

## Abstract

Neuronal ankyrins cluster and link membrane proteins to the actin and spectrin-based cytoskeleton. Among the three vertebrate ankyrins, little is known about neuronal Ankyrin-R (AnkR). We report AnkR is highly enriched in Pv^+^ fast-spiking interneurons in mouse and human. We identify AnkR-associated protein complexes including cytoskeletal proteins, cell adhesion molecules (CAMs), and perineuronal nets (PNNs). We show that loss of AnkR from forebrain interneurons reduces and disrupts PNNs, decreases anxiety-like behaviors, and changes the intrinsic excitability and firing properties of Pv^+^ fast-spiking interneurons. These changes are accompanied by a dramatic reduction in Kv3.1b K^+^ channels. We identify a novel AnkR-binding motif in Kv3.1b, and show that AnkR is both necessary and sufficient for Kv3.1b membrane localization in interneurons and at nodes of Ranvier. Thus, AnkR regulates Pv^+^ fast-spiking interneuron function by organizing ion channels, CAMs, and PNNs, and linking these to the underlying *β*1 spectrin-based cytoskeleton.

## INTRODUCTION

Ion channels and cell adhesion molecules (CAMs) are frequently recruited to, stabilized, and maintained at specific neuronal membrane domains by scaffolding proteins. The ankyrin scaffolding proteins, consisting of Ankyrin-R, -B, and -G (AnkR, AnkB, and AnkG, respectively), are the primary link between the submembranous spectrin-based cytoskeleton and the cytoplasmic domains of many transmembrane proteins (Michaely and Bennett, 1995; Sedgwick and Smerdon, 1999). For example, AnkG links Na^+^ and K^+^ channels, and the CAM neurofascin 186 (NF186) at axon initial segments (AIS) and nodes of Ranvier to the underlying *β*4 spectrin and actin-based cytoskeleton. The clustering of channels at the AIS and nodes facilitates fast and efficient action potential propagation (Dzhashiashvili et al., 2007; Zhou et al., 1998). Similarly, AnkB stabilizes Na^+^ channels and L1CAM family membrane proteins in unmyelinated axons and at paranodal junctions of myelinated axons by linking these membrane proteins to *β*2 spectrin (Chang et al., 2014; Scotland et al., 1998; Susuki et al., 2018). However, little is known about the function of AnkR in the nervous system. Instead, AnkR has mostly been studied in red blood cells where it maintains the cell’s structural integrity via its link between *β*1 spectrin and the cytoplasmic domain of the anion exchanger Band 3 (Bennett and Stenbuck, 1979). Loss of AnkR results in fragile erythrocyte membranes and hemolytic anemia (Lux et al., 1990). Intriguingly, case studies of patients with hereditary spherocytic anemia, caused by mutations in AnkR, report various neurological disturbances (Coetzer et al., 1988; McCann and Jacob, 1976; Miya et al., 2012), and a number of recent epigenome-wide association studies in Alzheimer’s disease (AD) have consistently found neuropathology-associated DNA hypermethylation of *ANK1* (*ANK1* is the gene encoding AnkR) (De Jager et al., 2014; Gasparoni et al., 2018; Higham et al., 2019; Lunnon et al., 2014; Smith et al., 2019a; Smith et al., 2019b). Additionally, AnkR can substitute for AnkG to cluster Na^+^ channels at nodes of Ranvier (Ho et al., 2014) but not AIS (Liu et al., 2020a). Thus, AnkR may play important, but as yet undefined, roles in nervous system function in both the healthy and diseased brain.

In addition to clustering ion channels, and through its interaction with CAMs, AnkG assembles and maintains a complex extracellular matrix (ECM) consisting of chondroitin sulfate proteoglycans (CSPGs) and other ECM proteins that surround AIS and nodes of Ranvier (Amor et al., 2017; Hedstrom et al., 2007; Susuki et al., 2013). Thus, ankyrins may function generally to link ECMs to the cytoskeleton through their membrane receptors. One highly condensed and specialized ECM in the nervous system is the perineuronal net (PNN). PNNs surround synaptic innervations and are thought to be important to maintain the balance of excitation and inhibition (Carceller et al., 2020). The majority of PNNs surround the soma and proximal dendrites of fast-spiking parvalbumin-positive (Pv^+^) inhibitory interneurons and have a CSPG composition similar to the perinodal and AIS ECM (Fawcett et al., 2019). However, how PNNs themselves are assembled, maintained, and restricted to specific domains and neuronal subtypes, is unknown.

Here, we show the loss of AnkR from GABAergic forebrain neurons results in a reduction and altered structure of PNNs, a reduction in anxiety-like behaviors, and altered intrinsic excitability and firing properties of PNN^+^ fast-spiking interneurons. We identify AnkR-interacting adhesion molecules that may tether PNNs to the spectrin cytoskeleton. Importantly, the altered excitability reflects the loss of Kv3.1b K^+^ channels. We identify the motif in Kv3.1b necessary for its interaction with AnkR. We show AnkR is both necessary and sufficient for the recruitment and clustering of Kv3.1b K^+^ channels in the neuronal membrane.

## RESULTS

### AnkR is highly enriched in Pv^+^ inhibitory interneurons

To determine where AnkR is located, we immunostained mouse forebrain using antibodies against AnkR. We found that AnkR is highly enriched in the soma and proximal dendrites of a subset of neurons sparsely distributed throughout the cortex and hippocampus (**Fig. 1A**). Immunoblotting of brain homogenates shows AnkR protein in the brain increases during early postnatal development, peaking by postnatal day 30 (**Fig. 1B**). The sparse distribution of AnkR-labeled cells was highly reminiscent of the distribution of cortical interneurons. Indeed, immunostaining with antibodies against parvalbumin (Pv), a marker of fast-spiking interneurons (**Fig. 1C, D**), shows >90% of Pv^+^ cells in postnatal day 56 (P56) cortex and hippocampus have high levels of AnkR, while only ∼70% of AnkR^+^ cells in cortex and hippocampus are Pv^+^. AnkR is also highly expressed in Pv^+^ neurons in human brain. AnkR staining was identified in laminae II-VI in human cortical biopsy samples. Labeling was strong and membranous and was most intense in larger neurons of laminae III and V (**Fig. 1E**). AnkR staining intensity in large neurons of lamina III and V was similar to staining intensity of erythrocytes (not shown). Furthermore, AnkR labeling was only identified in a subset of neurons in human cortex and staining was not present in endothelial cells or glia. A subset of AnkR^+^ neurons in laminae III and V co-expressed Pv in the neuronal cytoplasm (**Fig. 1E**), although, as in mice, the total number of AnkR^+^ positive neurons exceed the number of Pv^+^ cells. These results expand on those previously reported in rat neocortex (Wintergerst et al., 1996).

**Figure 1.**
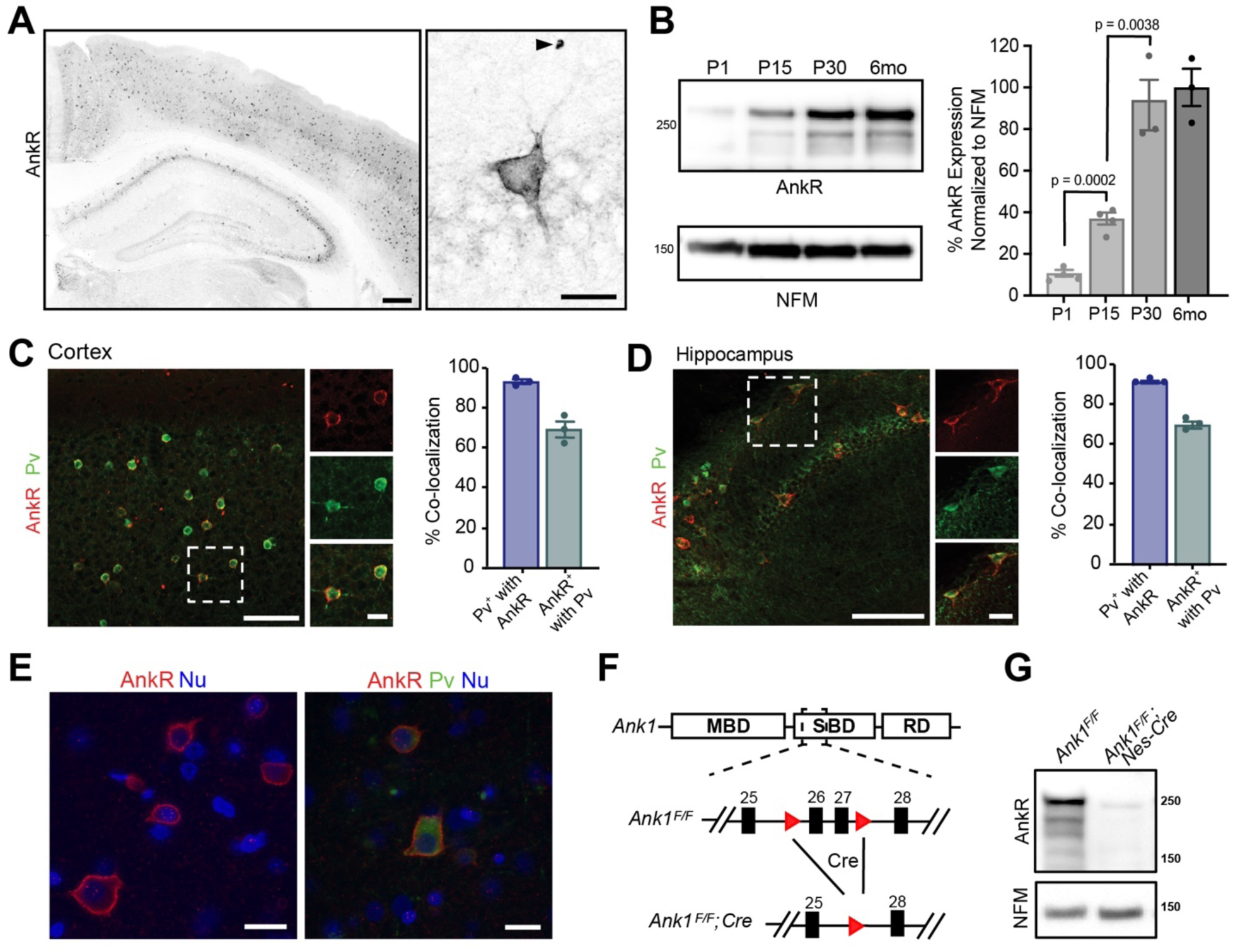
AnkR is expressed in select neurons in the cortex and hippocampus. (A) Immunostaining of coronal mouse brain for AnkR. Arrowhead indicates a red blood cell. Scalebars, left, 250 μm, right, 20 µm. (B) Immunoblot of P1, P15, P30, and 6-month C57BL/6 brain homogenates for anti-AnkR and neurofilament-M (NFM). Quantification of immunoblot in ImageJ by comparison of AnkR to NFM loading control from three independent duplicate experiments (n=3-4 mice/group). Samples were normalized to NFM, then compared to 6-month animals. (**C, D**) Immunostaining for AnkR (red) and Parvalbumin (Pv, green) in P56 coronal cortex (**C**, N = 3 mice; n=4495 cells) and coronal hippocampus (**D**, N = 3 mice; n=1637 cells). Scalebars, 50 µm; inset, 10 µm. (E) Immunostaining of human cortex for AnkR (red) and Parvalbumin (green). Nuclei are labeled using DAPI (blue). Scalebars, 20 µm. (F) Schematic of the *Ank1* conditional allele. *loxP* sites (red triangles) flank exons 26 and 27 in the spectrin binding domain (SBD), after the membrane binding domain (MBD) and before the regulatory domain (RD). Cre-mediated recombination and removal of exons 26 and 27 generates a premature stop codon in exon 28. (G) Immunoblot of brain homogenates for anti-AnkR, and Neurofilament-M. Molecular weights are indicated at right in kDa. All error bars indicate mean ± SEM.

To determine the role of neuronal AnkR and confirm its high expression in interneurons, we constructed a floxed allele for *Ank1* (*Ank1^F/F^*; **Fig. 1F**); this new model allows for an exploration of AnkR function in the brain while avoiding the confound of anemia due to loss of AnkR from red blood cells. We removed AnkR from the nervous system using *Nestin-Cre* mice (*Ank1^F/F^;Nes-Cre*), and from GABAergic forebrain interneurons using *Dlx5/6-Cre* mice (*Ank1^F/F^;Dlx5/6-Cre)*. We confirmed the efficient loss of AnkR after recombination by immunoblot of *Ank1^F/F^;Nes-Cre* brain homogenates (**Fig. 1G**). Moreover, immunostaining of forebrain sections from both *Ank1^F/F^;Nes-Cre* and *Ank1^F/F^;Dlx5/6-Cre* showed efficient and selective loss of AnkR in neurons; AnkR expression in erythrocytes was unaffected (**Supplemental Fig. 1A**). These results also highlight the specificity of our antibodies. Additionally, hemoglobin levels in *Ank1^F/F^;Nes-Cre* and *Ank1^F/F^;Dlx5/6-Cre* mice were normal (data not shown). Although most Pv^+^ cells are AnkR^+^ (**Fig. 1C, D**) there is a subpopulation of AnkR^+^ neurons that are not Pv^+^; these cells are most likely another subtype of GABAergic neuron since neurons with high levels of AnkR immunoreactivity in cortical and hippocampal neurons were not detected in *Ank1^F/F^;Dlx5/6-Cre* mice (**Supplemental Fig. 1A**). Interestingly, and in contrast to the rescue of AnkG by AnkR at nodes of Ranvier(Ho et al., 2014), we found no evidence for reciprocal compensation by AnkG in AnkR-deficient neurons. AnkG in Pv^+^ neurons in cortex remained highly restricted to the AIS and nodes in all genotypes analyzed (**Supplemental Fig. 1B** and data not shown). Together, these results show that AnkR is abundantly expressed in Pv^+^ interneurons of the forebrain, that its localization is distinct from that of AnkB and AnkG, and that we have generated a floxed *Ank1* allele that allows for cell-type specific deletion in the nervous system.

### Elucidating the AnkR interactome

What functions does AnkR have in GABAergic forebrain interneurons? To begin to answer this question we determined AnkR’s interactome. Since AnkR and *β*1 spectrin are binding partners in erythrocytes and can function together at nodes of Ranvier to stabilize Na^+^ channels (Liu et al., 2020b), we first showed that AnkR and *β*1 spectrin also form a protein complex in Pv^+^ interneurons. Immunostaining showed that *β*1 spectrin is highly expressed and colocalizes with AnkR in forebrain interneurons (**Fig. 2A**). Furthermore, *β*1 spectrin and AnkR reciprocally co-immunoprecipitate each other (**Fig. 2B**). Immunoblots of *Ank1^F/F^;Nes-Cre* mouse brain homogenates show that *β*1 spectrin protein levels are significantly reduced in the absence of AnkR compared to control mice (**Fig. 2C**). Similarly, immunostaining of cortex and hippocampus from *Ank1^F/F^;Nes-Cre* mice showed remarkably reduced *β*1 spectrin immunofluorescence compared to control mice (**Supplemental Fig. 2A, B**). Together, these results show that AnkR interacts with and maintains *β*1 spectrin in Pv^+^ forebrain interneurons.

**Figure 2.**
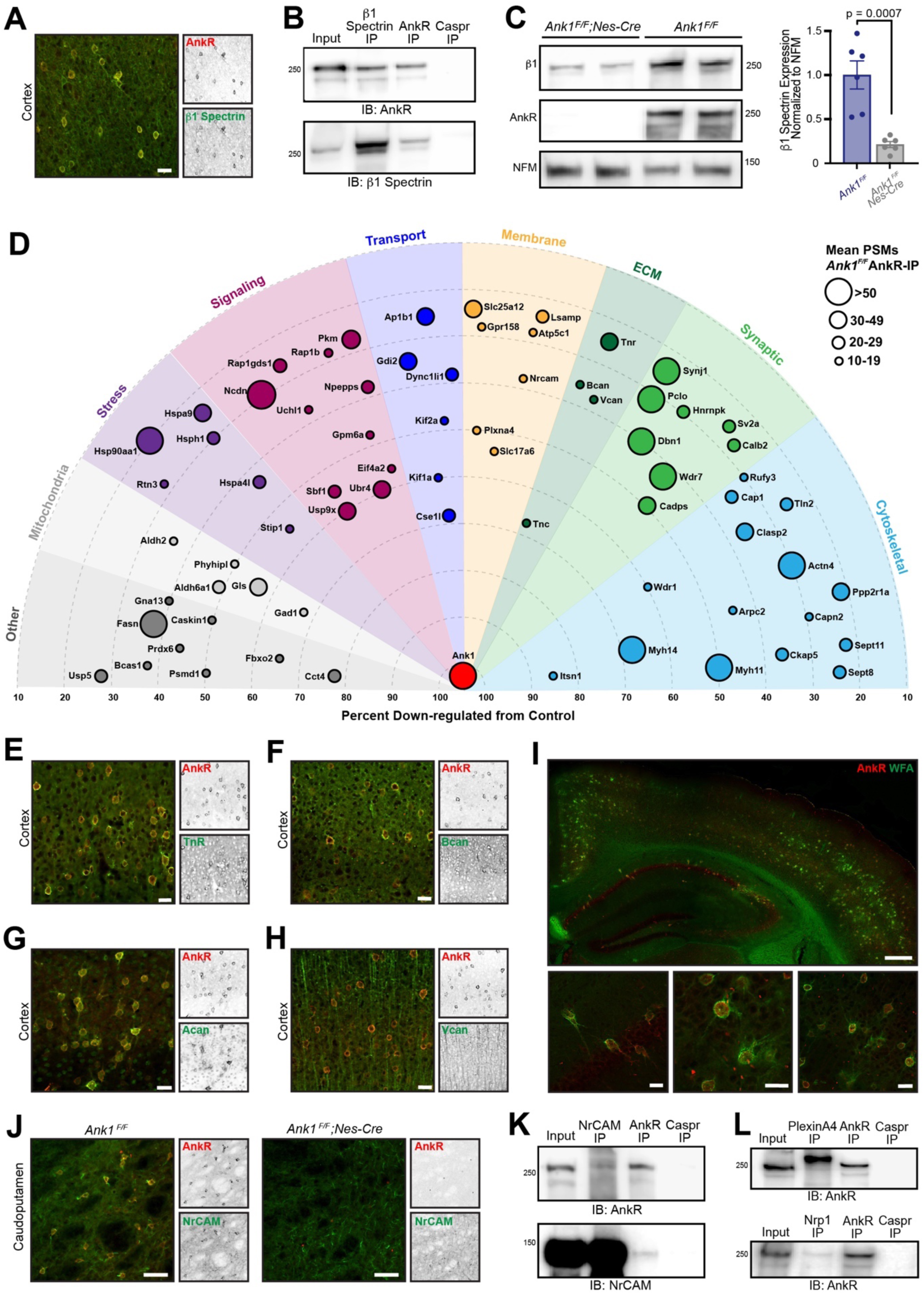
AnkR interacting proteins. (A) Immunostaining of mouse cortex using antibodies against AnkR (red) and *β*1 spectrin (green). Scalebar, 20 µm. (B) Immunoblot of *β*1 spectrin, AnkR, and Caspr immunoprecipitation reactions using antibodies against AnkR and *β*1 spectrin. IP, immunoprecipitation; IB, immunoblot. (C) Immunoblot of control and AnkR-deficient mouse brains using antibodies against *β*1 spectrin, AnkR, and neurofilament-M (NFM). Quantification of the *β*1 spectrin immunoblots normalized to NFM. Error bars indicate mean ± SEM. N=6. (D) Top AnkR-interacting candidates. Circle size corresponds to the mean PSM from IP mass spectrometry. Concentric circles radiating from *Ank1* correspond to % reduction in PSMs from AnkR knockout mouse compared to control. Identified proteins are organized according to their putative functions. (**E-H**) Immunostaining of mouse cortex using antibodies against AnkR (red) and TnR (green, **E**), Bcan (green, **F**), Acan (green, **G**), and Vcan (green, **H**). Scalebars, 20 μm. (I) Immunostaining of mouse cortex and hippocampus using antibodies against AnkR (red) and fluorescent-labeled WFA (green). Scalebars, 250 µm (top) and 20 µm (bottom). (J) Immunostaining of control (left) and AnkR-deficient mouse brain (right) using antibodies against AnkR (red) and NrCAM (green). Scalebars, 50 µm. (K) Immunoblot of NrCAM, AnkR, and Caspr immunoprecipitation reactions using antibodies against AnkR (top) and NrCAM (bottom). (L) Immunoblot of PlexinA4, AnkR, and Caspr immunoprecipitation reactions using antibodies against AnkR (top). Immunoblot of Nrp1, AnkR, and Caspr immunoprecipitation reactions using antibodies against AnkR (bottom).

To identify additional AnkR interacting proteins, we combined two unbiased mass spectrometry screens (**Supplemental Fig. 2C**). First, we performed mass spectrometry on AnkR immunoprecipitations (IPs) in biological triplicate from whole *Ank1^F/F^* mouse brain lysates. These yielded 3241 unique proteins with at least one peptide spectral match (PSM) in each sample. We further narrowed the number of potential AnkR-binding proteins by setting an arbitrary threshold of ≥10 mean PSMs found in the IPs. As a second, orthogonal approach, we performed differential proteomics and mass spectrometry using wildtype (*Ank1^+/+^* or WT) and AnkR-deficient (*Ank1 ^pale/pale^* or AnkR-KO)(Ho et al., 2014) hindbrain homogenates (**Supplemental Fig. 2C**). We used AnkR-KO mice rather than *Ank1^F/F^;Nes-Cre* mice to avoid confounds due to incomplete recombination or contributions from cells still expressing AnkR. Mass spectrometry yielded 2465 unique proteins, of which 986 were reduced in AnkR-KO mice compared to WT. Since ankyrins function as scaffolds that stabilize and retain membrane proteins, we reasoned that loss of AnkR might result in increased turnover and lower amounts of AnkR-interacting proteins. Therefore, we focused only on those proteins that had fewer PSMs in the AnkR-KO compared to WT. To further refine our analysis, we set an arbitrary threshold such that potential AnkR-interacting proteins must have ≥20% reduction in PSMs, with ≥5 PSMs found in WT mice and a ≥3 PSMs difference between WT and AnkR-KO. Combining the two data sets revealed 72 potential AnkR-interacting proteins (**Supplemental Fig. 2C**).

We sorted these 72 proteins into functional categories and plotted them using concentric rings to indicate the percent reduction in AnkR-KO mice, with circle size representing the mean number of PSMs in the IPs (**Fig. 2D**). Among the proteins enriched and passing our stringent filtering criteria, we found cytoskeletal, membrane, signaling, and ECM proteins. Surprisingly, although AnkR is a cytoplasmic scaffolding protein, the ECM proteins TenascinC (TnC), TenascinR (TnR), Brevican (Bcan) and Versican (Vcan) were all identified in the AnkR IPs and also enriched in WT compared to AnkR-KO. We previously reported that AnkG, through the CAM NF186, interacts with and recruits the CSPGs Bcan and Vcan to AIS and nodes of Ranvier (Hedstrom et al., 2008; Susuki et al., 2013). TnR is also found at nodes and binds to Vcan and Bcan (Bekku et al., 2009). Immunostaining of cortex revealed that antibodies against TnR and Bcan strongly label AnkR^+^ neurons (**Figs. 2E, F**); Vcan immunoreactivity was not restricted to AnkR^+^ neurons and was more widely distributed (**Fig. 2H**). TnR, Bcan and Vcan are well known components of PNNs (Fawcett et al., 2019; Wintergerst et al., 1996). Immunostaining for the CSPG and PNN protein Aggrecan (Acan) (Carulli et al., 2007; Rowlands et al., 2018) also showed strong colocalization with AnkR (**Fig. 2G**); although Acan was detected by mass spectrometry, it did not pass our stringent filtering. In addition to antibodies against Bcan, TnR, and Acan, PNNs can also be detected using the fluorescently-labeled *Wisteria Floribunda* (WFA) lectin (Bruckner et al., 1993), which binds to *N*-acetyl-D-glucosamine at the ends of chondroitin sulfate chains. Co-staining of WFA and AnkR shows that AnkR^+^ neurons are surrounded by WFA-labeled PNNs (**Fig. 2I**).

How can AnkR, an intracellular scaffolding protein, interact with extracellular PNNs? We reasoned this could occur through CAMs that bridge AnkR and PNNs. Our list of potential AnkR-binding CAMs included two strong candidates: NrCAM and PlexinA4. NrCAM is a member of the L1 family of CAMs with known ankyrin-binding activity (Davis and Bennett, 1994). However, NrCAM can also be shed from the cell surface and incorporated into the ECM surrounding nodes of Ranvier through direct binding to NF186 (Susuki et al., 2013). Thus, NrCAM can function both as a membrane receptor and as a component of the perinodal ECM. Immunostaining for NrCAM showed strong colocalization with both AnkR^+^ and WFA^+^ neurons in caudoputamen (**Fig. 2J** and **Supplemental Fig. 2G**), but less robust colocalization in hippocampus and cortex (**Supplemental Figs. 2D-F**). These results suggest AnkR may function together with NrCAM in a subset of GABAergic neurons and emphasizes the diversity of PNNs and their interacting proteins. Somatodendritic, but not AIS, NrCAM immunoreactivity was dramatically reduced in *Ank1^F/F^;Nes-Cre* mouse brain (**Fig. 2J**). Furthermore, NrCAM and AnkR reciprocally co-immunoprecipitate each other, while the axonal membrane protein Caspr, normally found at paranodal junctions of myelinated axons, does not (**Fig. 2K**). These results support recent work exploring the molecular heterogeneity of PNNs and suggests another potential PNN subtype involving a unique CAM membrane receptor (Irvine and Kwok, 2018; Yamada and Jinno, 2017).

PlexinA4 functions together with Neuropilin-1 (Nrp1) as a receptor for semaphorin signaling (Nakamura et al., 2000). Sema3A is a component of the PNNs surrounding Pv^+^ interneurons (Kwok et al., 2011); moreover, enzymatic or genetic disruption of PNNs reduces Sema3A (de Winter et al., 2016). PlexinA4 and Nrp1 are widely expressed throughout the nervous system. Although immunostaining did not reveal any specific enrichment for PlexinA4 in AnkR^+^ interneurons, immunostaining for Nrp1 showed strong enrichment in AnkR^+^ neurons in deep cerebellar nuclei (**Supplemental Fig. 2I**). Nevertheless, we found that PlexinA4 and Nrp1 co-immunoprecipitated with AnkR from brain homogenates (**Fig. 2L**). Together, our proteomic studies show that AnkR co-localizes with multiple PNN proteins and may indirectly interact with PNNs through the membrane receptors NrCAM and PlexinA4. Other membrane proteins identified in our proteomics, but not analyzed here, may also function to link PNNs to AnkR. Based on these proteomic, biochemical, and immunostaining results, we focused on the relationship between AnkR and PNNs.

### AnkR is required to maintain PNN density and structure

To determine if AnkR contributes to the formation, maintenance, and structure of PNNs, we used WFA to label PNNs in cortex and hippocampus of one-month-old *Ank1^F/F^*, *Ank1^+/+^;Dlx5/6-Cre*, and *Ank1^F/F^;Dlx5/6-Cre* mice (**Fig. 3A**). At this age, we found little difference in the number of WFA^+^/Pv^+^, WFA^-^/Pv^+^, or WFA^+^/Pv^-^ neurons per unit area (UA) in either hippocampus or cortex (**Fig. 3B**). However, when we measured the fluorescence intensity of WFA, we found a significant reduction in both cortex and hippocampus in *Ank1^F/F^;Dlx5/6-Cre* mice compared to floxed or Cre controls (**Fig. 3C**); further subdividing cortical regions showed a similar decrease (**Supplemental Figs. 3A, C**). Examination of PNNs at high magnification showed that in the absence of AnkR, PNNs were less compact compared to control mice. We classified the PNNs as being dense (0), having a few small holes (1), or having large numerous holes (2) (**Supplemental Fig. 3D**). AnkR-deficient neurons were significantly less likely to be compact and more likely to have holes in their nets (**Figs. 3D, E**). In 12-month-old mice we found a significant reduction in the number of WFA^+^/Pv^+^ cells per unit area in the hippocampus and cortex of *Ank1^F/F^;Dlx5/6-Cre* mice compared to floxed or Cre controls (**Figs. 3F, G**). In addition, and like in the one-month-old *Ank1^F/F^;Dlx5/6-Cre* mice, we measured a ∼50% decrease in the WFA fluorescence intensity in both hippocampus and cortex (**Fig. 3H**). This significant reduction in WFA was seen in all cortical regions (**Supplemental Fig. 3B**). The normally compact PNN structure, as observed in 12-month-old control mice, was also significantly disrupted in *Ank1^F/F^;Dlx5/6-Cre* mice with even more prominent holes in the PNNs compared to either control or one-month-old *Ank1^F/F^;Dlx5/6-Cre* mice (**Figs. 3I, J, and Supplemental Movies 1, 2**). Furthermore, the strong reduction in NrCAM immunoreactivity seen in *Ank1^F/F^;Nes-Cre* mouse brain (**Fig. 2J**) was matched by a strong reduction in WFA^+^ cells in the same region (**Supplemental Figs. 2G, H**). Together, these results suggest that loss of AnkR does not disrupt the ability of PNNs to form, but rather AnkR helps maintain PNNs and their normal compact structure through binding to PNN-interacting CAMs like NrCAM and PlexinA4.

**Figure 3.**
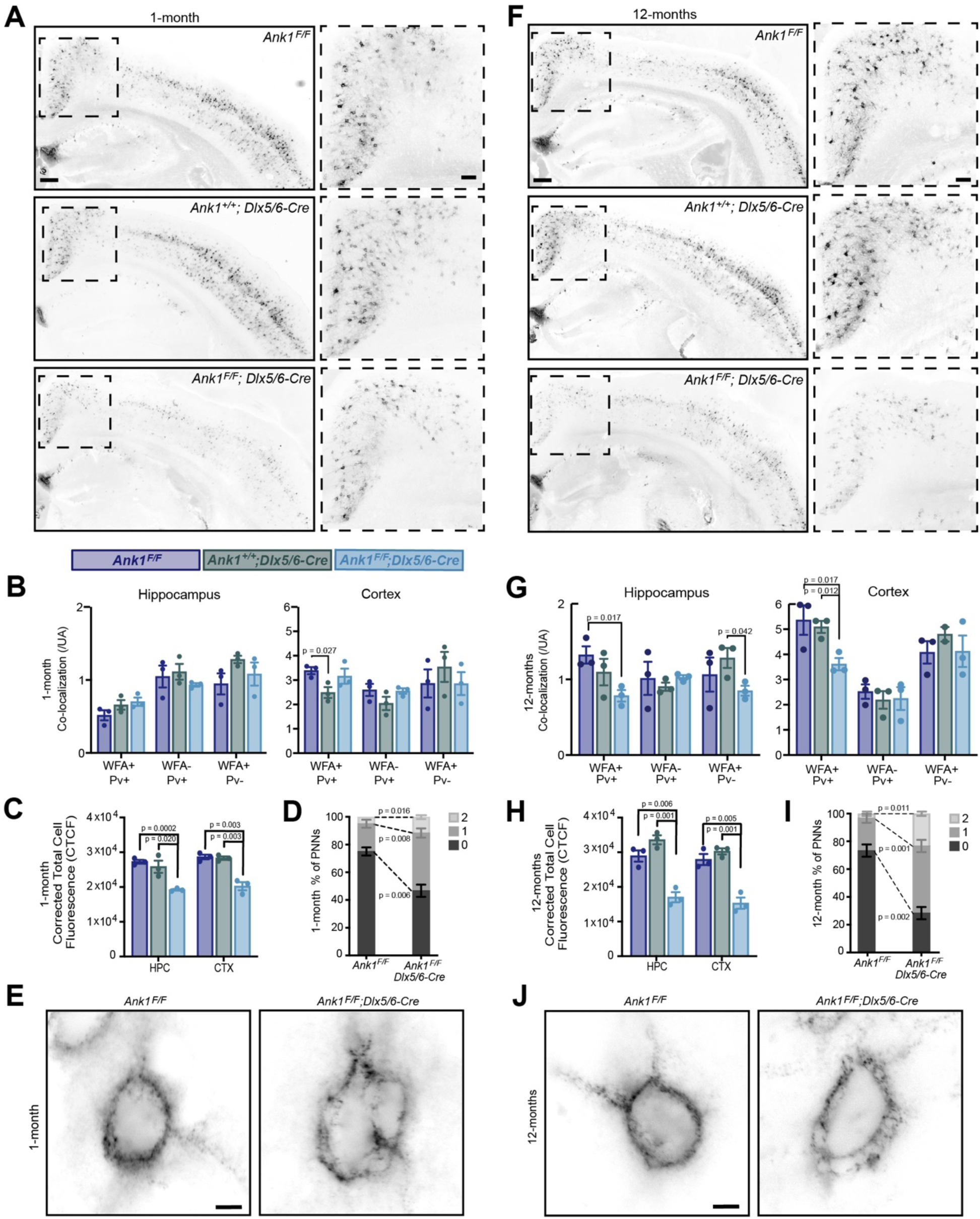
AnkR maintains perineuronal net density and structure. (A) Fluorescent WFA labeling of PNNs in coronal sections of cortex and hippocampus in one-month-old mice. Genotypes of the respective mice are indicated. Boxed regions are shown to the right. Scalebars, 250 µm and 100 µm. (B) Quantification of colocalization between WFA and Pv labeling per unit area (UA) in hippocampus and cortex. N=3 mice/group. (C) Corrected total cell fluorescence (CTCF) in hippocampus (HPC) and cortex (CTX). N=3 mice/group, n=230 cells/animal. (D) Semi-quantitative analysis of high magnification WFA showing increased hole size and disruption of PNNs in retrosplenial (RSP) cortex of 1-month mice. N=3 mice/group, n=20 cells/animal. (E) Fluorescent WFA label of PNNs in RSP cortex of one-month-old mice. Genotypes of the respective mice are indicated. Scalebars, 5 µm. (F) Fluorescent WFA labeling of PNNs in coronal sections of cortex and hippocampus in 12-month-old mice. Genotypes of the respective mice are indicated. Boxed regions are shown to the right. Scalebars, 250 µm and 100 µm. (G) Quantification of colocalization between WFA and Pv labeling per unit area (UA) in hippocampus and cortex. N=3 mice/group. (H) CTCF in HPC and CTX. N=3 mice/group, n=230 cells/animal. (I) Semi-quantitative analysis of high magnification WFA showing increased hole size and disruption of PNNs in RSP cortex of 12-month mice. N=3 mice/group, n=20 cells/animal. (J) Fluorescent WFA label of PNNs in RSP cortex of 12-month-old mice. Genotypes of the respective mice are indicated. Scalebars, 5 µm. Error bars indicate mean ± SEM. N=3/group.

### Loss of AnkR from GABAergic forebrain interneurons decreases anxiety-like behaviors

To determine if loss of AnkR from GABAergic interneurons alters nervous system function, we analyzed the behavior of *Ank1^F/F^;Dlx5/6-Cre* mice compared to control mice (*Ank1^F/^*^F^ and *Ank1^+/+^;Dlx5/6-Cre*). We first performed a 30-minute open field assessment to rule out locomotor deficits (**Fig. 4A** and **Supplemental Fig. 4A-C**) since a hypomorph of AnkR was previously reported to have a loss of Purkinje neurons by 6 months of age (Peters et al., 1991). We found that all genotypes analyzed had normal locomotion, but both *Ank1^+/+^;Dlx5/6-Cre* and *Ank1^F/F^;Dlx5/6-Cre* mice showed increased velocity and distance traveled compared to *Ank1^F/^*^F^ mice. These results are consistent with previous reports that the *Dlx5/6-Cre* transgene results in a hyperactive phenotype characterized by increased velocity and movement(de Lombares et al., 2019). Nevertheless, during the open field assessment, *Ank1^F/F^;Dlx5/6-Cre* mice spent significantly more time in the center of the arena and less time in the perimeter compared to both *Ank1^F/^*^F^ and *Ank1^+/+^;Dlx5/6-Cre* mice (**Fig. 4B, C**), suggesting that loss of AnkR may be anxiolytic.

**Figure 4.**
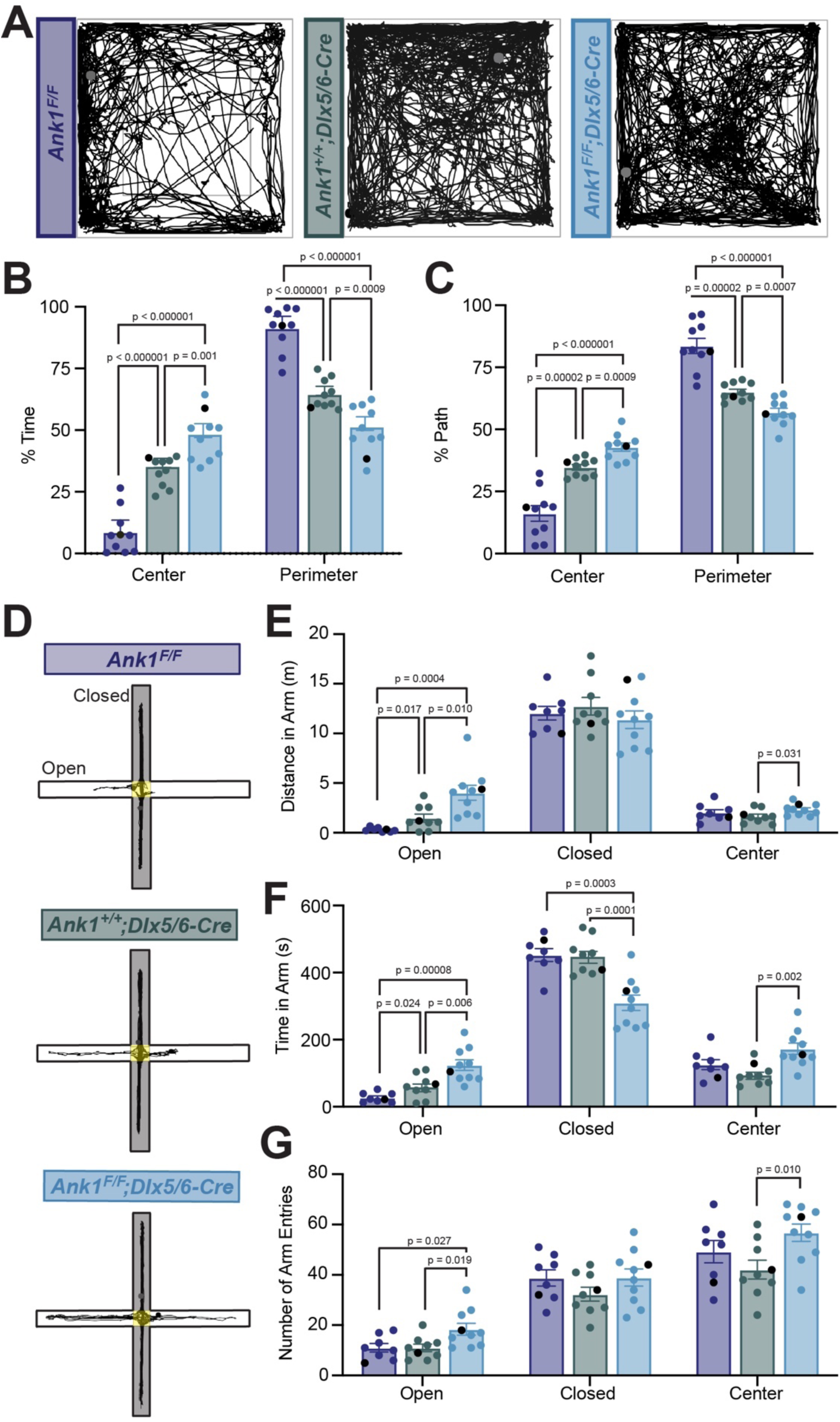
Mice lacking AnkR in GABAergic forebrain interneurons have reduced anxiety. (A) 30 minute-long recordings of mouse trajectories in the open field assay. Genotypes are indicated. (B) The percent of time spent in the center or perimeter of the open field. (C) The percent of the total path spent in the center or perimeter of the open field. (D) 10 minute long recordings of mouse trajectories in the elevated plus maze. (E) The distance traveled in the open arm, closed arm, or the center of the elevated plus maze. (F) The time spent in the open arm, closed arm, or the center of the elevated plus maze. (G) The number of entries into the open arm, closed arm, or center of the elevated plus maze. In all panels error bars indicate mean ± SEM. Black circles indicate the animals corresponding to the representative traces.

To further distinguish between increased activity and decreased anxiety-like behaviors, we used the elevated plus maze (EPM) (**Fig. 4D**). In contrast to the open field assessment, during a 10-minute trial in the EPM we observed no difference among the three genotypes in the velocity or total distance traveled in all arms (**Supplemental Fig. 4D-E**). However, we measured a significant increase in the distance traveled in the open arms, the time spent in the open arms, and the number of open arm entries performed by *Ank1^F/F^;Dlx5/6-Cre* mice compared to control mice (**Fig. 4E-G**). Together, these data suggest that loss of AnkR from forebrain GABAergic neurons reduces anxiety-like behaviors.

### AnkR influences the intrinsic excitability and firing properties of WFA^+^ fast-spiking interneurons without altering their synaptic inputs

To determine how loss of AnkR impacts the intrinsic excitability and firing properties of fast-spiking interneurons, we performed whole-cell current clamp recordings on layer 5 PNN^+^ interneurons in somatosensory cortical slices from *Ank1^F/F^*, *Ank1^+/+^;Dlx5/6-Cre*, and *Ank1^F/F^;Dlx5/6-Cre* mice. Fast-spiking interneurons were identified in live brain slices by labeling PNNs with fluorescent WFA (**Supplemental Fig. 5A-C**). Indeed, 64 out of 66 WFA^+^ cells recorded were fast-spiking interneurons and the 2 non-fast spiking cells were not included in the analysis. We found that the resting membrane potential, input resistance, membrane capacitance, and rheobase current were not significantly altered in *Ank1^F/F^;Dlx5/6-Cre* neurons compared to controls (**Supplemental Table 1**). We determined the action potential latency, threshold, amplitude, half-width, afterhyperpolarization (AHP) amplitude, and AHP time from the single action potential elicited by the rheobase current (**Figure 5A**). Loss of AnkR decreased the action potential latency and threshold (**Fig. 5B, C**, and **Supplemental Table 1**) without changing the amplitude of the action potential (**Fig. 5D**, and **Table 1**). Interestingly, loss of AnkR also altered the shape of the action potential, resulting in a 47% broader action potential with a shallower and delayed AHP (**Fig. 5E-H,** and **Supplemental Table 1**).

**Figure 5.**
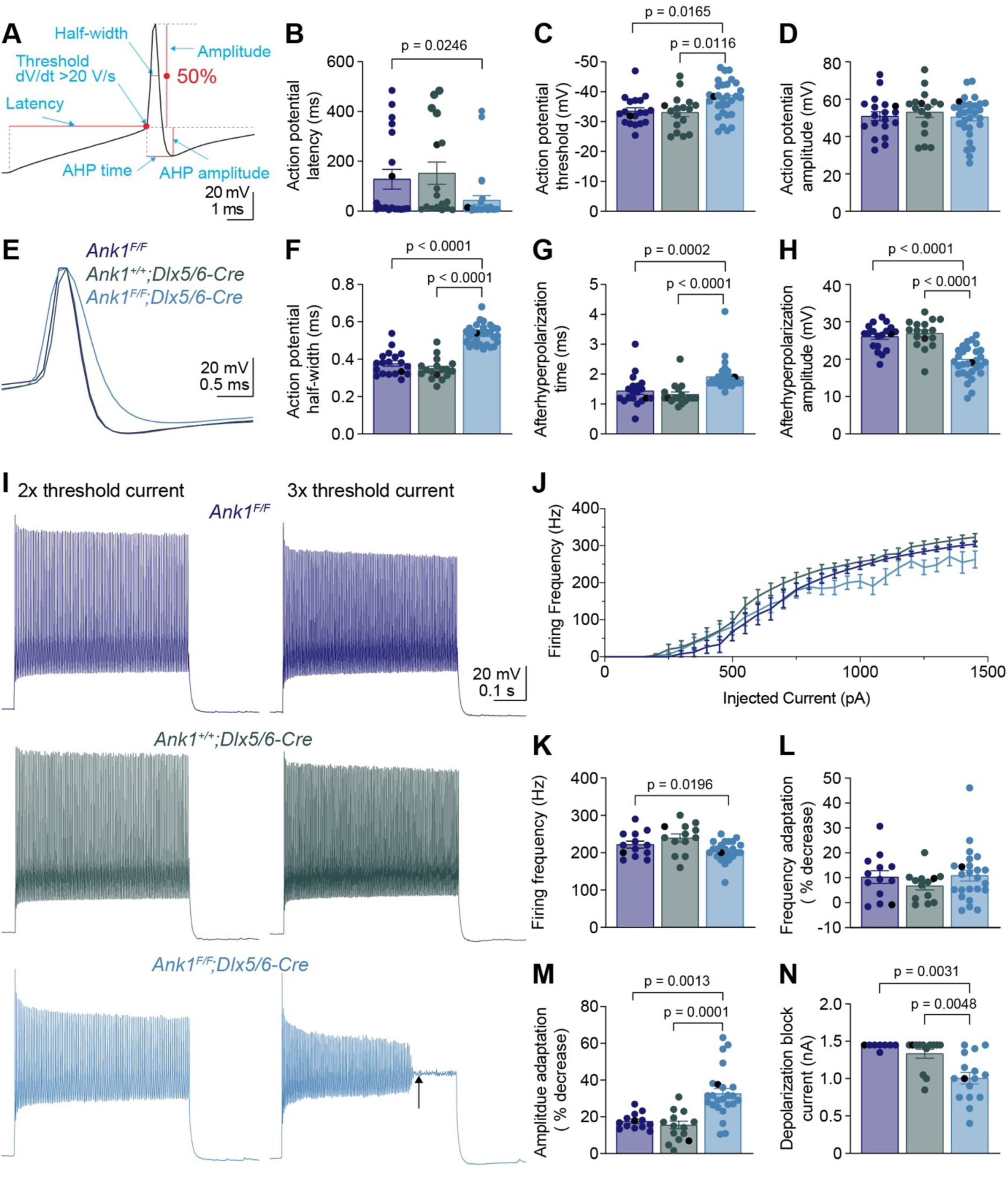
WFA^+^ neurons in *Ank1^F/F^;Dlx5/6-Cre* mice have abnormal action potentials and spike train characteristics. (**A**) A representative action potential illustrating the measurement of action potential parameters. AHP, afterhyperpolarization. (**B-D**) Summary data showing the action potential latency (**B**), action potential threshold (**C**), and action potential amplitude (**D**) from the single action potential evoked by rheobase current. (**E**) Representative single action potentials evoked by rheobase currents from *Ank1^F/F^*, *Ank1^+/+^;Dlx5/6-Cre*, and *Ank1^F/F^;Dlx5/6-Cre* cells. Action potentials are aligned at 50% of the rising phase on X axis and peak on Y axis. Note the wider action potential with shallower and delayed afterhyperpolarization in *Ank1^F/F^;Dlx5/6-Cre* cell. (**F-H**) Summary data showing the action potential half-width (**F**), afterhyperpolarization time (**G**), and afterhyperpolarization amplitude (**H**) from the single action potentials evoked by rheobase currents. (I) Representative spike trains from *Ank1^f/f^*, *Ank1^+/+^;Dlx5/6-Cre*, and *Ank1^F/F^;Dlx5/6-Cre* cells in response to 500-ms current injection. Left and right traces show the spike trains evoked by currents that are 2 and 3 times of the action potential threshold currents, respectively. Note the strong amplitude adaptation and premature depolarization block indicated by the arrow in the right trace of *Ank1^F/F^;Dlx5/6-Cre* cell. (J) The average firing frequency during 500-ms current injection as a function of injected currents. Note, recording was stopped at maximal 1450 pA current or when cells reached depolarization block. Since 13 out of 15 *Ank1^f/f^;Dlx5/6-Cre+* cells reached depolarization block prior to 1450 pA current while only 3 out of 19 control cells reached depolarization block prior to 1450 pA, the firing frequency is overestimated in the high current range for *Ank1^F/F^;Dlx5/6-Cre* neurons. (**K-M**) Summary data showing the average firing frequency during the first 100 ms (**K**), spike frequency adaptation (**L**), and amplitude adaptation (**M**) from the spike trains evoked by currents that are 2 times of the action potential threshold currents. (**N**) Summary data showing the minimal currents that caused the cells to enter depolarization block. If the maximal injected current (1450 pA) did not cause depolarization block, then 1450 pA was recorded as the result. For all panels, each circle represents one neuron and the black circles indicate the representative cells in (**E** and **I**). Bar graphs represent mean ± SEM. Statistical significance was determined by one-way ANOVA or Kruskal-Wallis test with multiple comparisons.

We also recorded trains of action potentials evoked by different levels of current injection (**Fig. 5I, J**). At 2 times the action potential threshold current, fast-spiking interneurons from *Ank1^F/F^;Dlx5/6-Cre* mice display a decreased firing frequency during the first 100 ms of current injection (**Fig. 5K**), but normal spike frequency adaptation (**Fig. 5L**). However, the spike amplitude adaptation was enhanced, resulting in a strong reduction in the amplitudes of action potentials towards the end of spike trains (**Fig. 5I, M**). In fact, *Ank1^F/F^;Dlx5/6-Cre* cells often entered depolarization block at much lower current levels than control cells (**Fig. 5I, N**). Thus, the reduction in firing frequency for *Ank1^F/F^;Dlx5/6-Cre* cells at high levels of current injection is actually an underestimation, because many *Ank1^F/F^;Dlx5/6-Cre* cells prematurely reached depolarization block and were not represented by the current-firing frequency curve (**Fig. 5J**).

In addition to examining intrinsic excitability of fast-spiking interneurons we also performed whole-cell voltage clamp recordings to record miniature postsynaptic currents. We observed no significant differences between the controls and *Ank1^F/F^;Dlx5/6-Cre* mice for the frequency and amplitude of miniature excitatory postsynaptic currents (mEPSC) or miniature inhibitory postsynaptic currents (mIPSC), as well as the ratio of excitatory to inhibitory inputs (E/I ratio) (**Supplemental Fig. 5D-I**). Taken together, these results show that loss of AnkR alters the intrinsic properties of WFA^+^ fast-spiking inhibitory interneurons. Furthermore, they are consistent with what has been reported in another model where PNNs of fast-spiking interneurons are disrupted due to loss of Bcan (Favuzzi et al., 2017); however loss of Bcan also induced changes in mPSCs. Intriguingly, the results are very similar to those seen in mice with loss of Kv3 K^+^ channel expression in fast-spiking interneurons (Lau et al., 2000). Hence, we hypothesize that loss of AnkR may impact Kv3 K^+^ channel function.

### AnkR recruits and maintains Kv3.1b K^+^ channels at the neuronal membrane

Among the Kv channels, Kv3.1b is highly expressed in WFA^+^ cortical interneurons (Hartig et al., 1999); Kv3.1b is also found at some CNS nodes of Ranvier, but not AIS (Devaux et al., 2003). Similarly, AnkR is enriched in WFA^+^ cortical interneurons (**Fig. 2I**), and can be found at some nodes of Ranvier (Ho et al., 2014), but not AIS (Liu et al., 2020a). Based on these similarities, we considered Kv3.1b to be a good candidate to interact with AnkR in somatic membranes of WFA^+^ cortical interneurons. Immunostaining of control 1- and 12-month-old somatosensory cortex showed that AnkR^+^ neurons were also Kv3.1b^+^ (**Fig. 6A,** and **Supplementary Fig. 6A**), and that AnkR and Kv3.1b colocalize at the neuronal membrane (**Fig. 6B**). This same colocalization was also seen in human cortex (**Fig. 6C**). Remarkably, AnkR-deficient *Ank1^F/F^;Dlx5/6-Cre* mice have a profound reduction in Kv3.1b^+^ neurons at both 1- and 12-months of age (**Figs. 6A, B**, and **Supplementary Fig. 6A**) and nearly complete loss of Kv3.1b immunofluorescence (**Fig. 6D**). Compared to controls, *Ank1^F/F^;Dlx5/6-Cre* mice have a ∼50% reduction in Kv3.1b protein (**Figs. 6E, F**), which persists in 12-month-old mice (**Supplementary Fig. 6B, C**). These results show that AnkR is required to maintain clustering of Kv3.1b in the somatic membrane of GABAergic interneurons.

**Figure 6.**
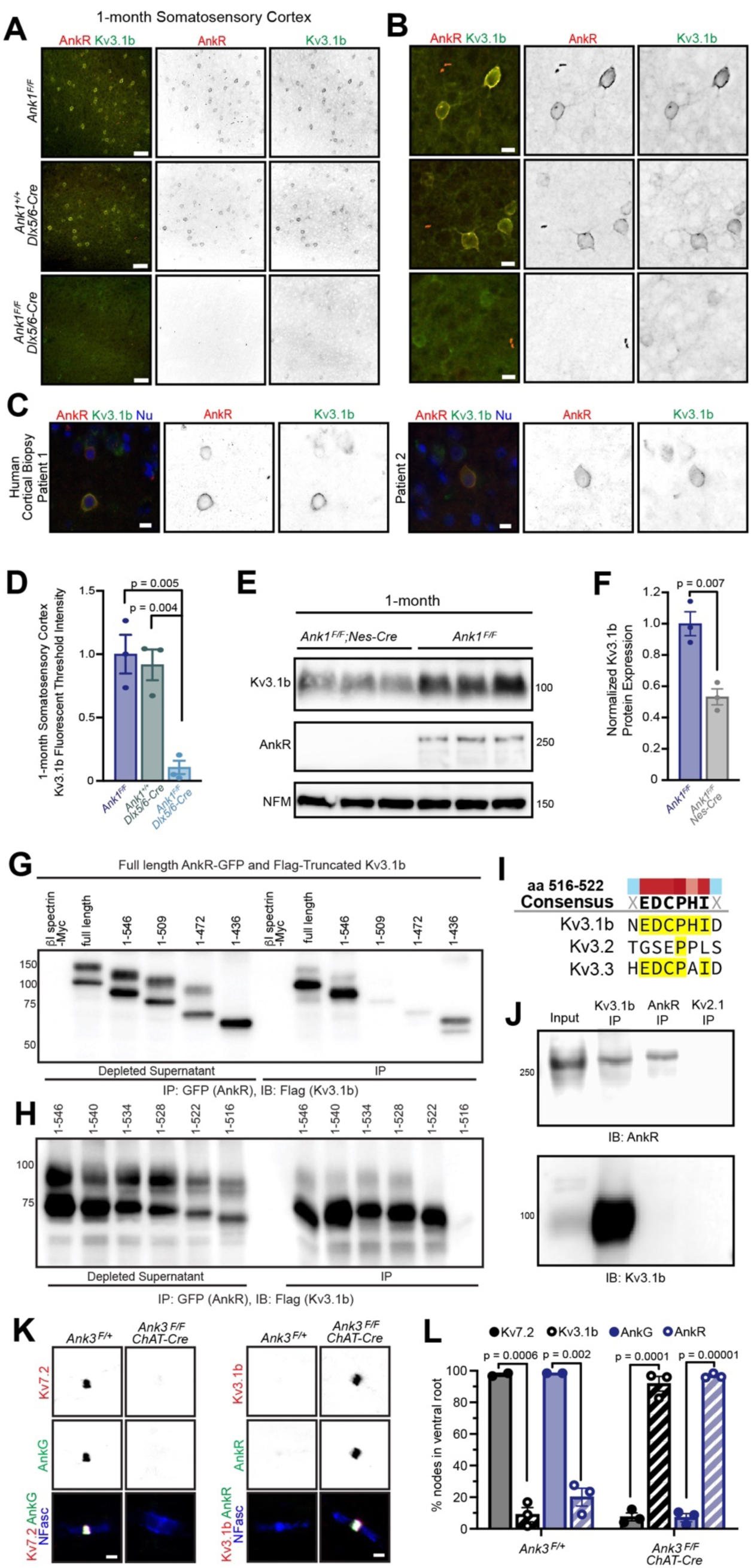
AnkR binds to Kv3.1b K^+^ channels and is both necessary and sufficient for its membrane localization and clustering. (**A, B**) Immunostaining of 1-month-old somatosensory cortex for AnkR (red) and Kv3.1b (green). Low magnification images are shown in (**A**) and high magnification images in (**B**). The genotypes analyzed are shown. Scalebars, 50 µm in (**A**) and 10 µm in (**B**). (C) Immunostaining of human cortical biopsies from two separate patients using antibodies against AnkR (red) and Kv3.1b (green), and DAPI (blue) to label nuclei (Nu). Scalebars, 10 µm. (D) Quantification of Kv3.1b immunofluorescence intensity in control and *Ank1^F/F^;Dlx5/6-Cre* mice. (E) Immunoblots of brain homogenates from 3 one-month-old control and 3 one-month-old AnkR-deficient brains using antibodies against Kv3.1b, AnkR, and NFM. (F) Quantification of Kv3.1b protein normalized to NFM. (**G, H**). Immunoblots of AnkR-GFP immunoprecipitations in cells co-expressing AnkR-GFP with Myc-tagged *β*1 spectrin, full length Flag-tagged Kv3.1b, or truncated versions of Flag-tagged Kv3.1b. The amino acids included in the Flag-tagged Kv3.1b truncation mutants are indicated. (I) The consensus AnkR-binding motif present in Kv3.1b and Kv3.3, but not Kv3.2. (J) Immunoblots of Kv3.1b, AnkR, and Kv2.1 immunoprecipitation reactions using antibodies against AnkR and Kv3.1b. (K) Immunostaining of ventral root nodes of Ranvier in *Ank3^F/+^ and Ank3^F/F^;ChAT-Cre* mice using antibodies against AnkG (green), Kv7.2 (red), and neurofascin (NFasc, blue) on the left, and AnkR (green), Kv3.1b (red), and NFasc (blue) on the right. Scalebars, 1μm. (L) Quantification of the percentage of nodes of Ranvier labeled for Kv7.2, Kv3.1b, AnkG, and AnkR in *Ank3^F/+^* and *Ank3^F/F^;ChAT-Cre* mice. 60-116 nodes/group. Error bars indicate mean ± SEM.

To determine if AnkR and Kv3.1b interact, we co-transfected HEK cells with AnkR-GFP and Flag-tagged Kv3.1b. AnkR-GFP efficiently co-precipitates full-length Flag-Kv3.1b (**Fig. 6G**). We then constructed serial C-terminal truncations of Flag-Kv3.1b. We found that AnkR-GFP pulled down amino acids (aa) 1-546 of Kv3.1b (**Fig. 6G**). However, additional shortening of the C-terminus blocked the interaction with AnkR. Thus, AnkR binds to the region of Kv3.1b including aa 510-546 (**Fig. 6G**). To further define the motif in Kv3.1b that interacts with AnkR, we generated additional C-terminal deletions of just 6 aa each, spanning aa 510-546 of Kv3.1b. We found that aa 516-522 of Kv3.1b (EDCPHI) are required for AnkR binding (**Fig. 6H**). A nearly identical motif is also present in Kv3.3, but not Kv3.2 (**Fig. 6I**). Kv3.3 is also highly expressed in Pv^+^ neurons (Chang et al., 2007) and immunostaining of control one-month-old somatosensory cortex showed that a subset of AnkR^+^ neurons also express Kv3.3 (**Supplementary Fig. 6D**). Using brain homogenates, we found that Kv3.1b, but not Kv2.1, co-immunoprecipitates AnkR (**Fig. 6J**). However, we were unable to detect Kv3.1b after immunoprecipitation of AnkR; this may suggest that only a small fraction of the total AnkR interacts with Kv3.1b. Together, these results show that AnkR binds directly to Kv3.1b.

Since Kv3.1b interacts with AnkR and is required for its membrane localization (**Figs. 6A, b**), we next determined if AnkR is sufficient to recruit Kv3.1b to neuronal membrane domains. Although some CNS nodes of Ranvier have clustered Kv3.1b, most PNS nodes of Ranvier normally have high levels of Kv7.2/3 K^+^ channels rather than Kv3.1b (**Figs. 6K, L**) (Pan et al., 2006). Kv7.2 K^+^ channel clustering requires binding to AnkG since Kv7.2 is absent from nodes in the ventral roots of AnkG-deficient (*Ank3^F/F^;ChAT-Cre*) mice (**Figs 6K, L**). AnkR clusters nodal Na^+^ channels(Ho et al., 2014) and NF186 in the ventral roots of *Ank3^F/F^;ChAT-Cre* mice (**Fig. 6K**). Although very few nodes in ventral root normally have Kv3.1b (**Figs. 6L**), the replacement of AnkG by AnkR in *Ank3^F/F^;ChAT-Cre* mice is sufficient to recruit and cluster Kv3.1b to nearly all nodes (**Figs. 6K, L**); and nodes in control spinal cord that have high levels of AnkR also have clustered Kv3.1b (**Supplemental Fig. 6E**). Similarly, AnkR recruits Kv3.3 K^+^ channels to nodes in AnkG-deficient ventral root axons (**Supplemental Fig. 6E**). Together, these results show that although AnkG and AnkR can both cluster Na^+^ channels at nodes of Ranvier, AnkG preferentially clusters Kv7.2/3 K^+^ channels and links them to the cytoskeleton through *α*2/*β*4 spectrin (Huang et al., 2017b), while AnkR is both necessary and sufficient to recruit Kv3.1b/3 K^+^ channels to neuronal membranes and nodes of Ranvier, and links them to the cytoskeleton through *α*2/*β*1 spectrin (Ho et al., 2014; Huang et al., 2017a) (**Supplemental Fig. 6G**). Thus, the type of K^+^ channel found at nodes of Ranvier is dictated by ankyrins.

## DISCUSSION

Ankyrins are well-known to function in neurons as scaffolds that link ion channels and membrane proteins to the spectrin cytoskeleton (Bennett and Lorenzo, 2013). We previously showed that in the absence of AnkG, AnkR can function at nodes of Ranvier as a secondary Na^+^ channel clustering mechanism (Ho et al., 2014). The rescue depends on AnkR’s recruitment to nodes from a pre-existing, unclustered pool. These findings motivated us to determine AnkR’s normal functions in the nervous system since it is unlikely that AnkR functions only as a backup for nodal Na^+^ channel clustering, since pathogenic *ANK1* variants are associated with nervous system dysfunction, and since altered methylation of *ANK1* is associated with AD. Our experiments confirm that in general, AnkR acts as a scaffolding protein like AnkB and AnkG. However, unlike AnkB and AnkG which are broadly expressed in all neurons, AnkR is highly and specifically enriched in subsets of neurons, including fast-spiking GABAergic interneurons, where it assembles and stabilizes unique protein complexes necessary for the proper function of these cells (**Fig. 7**).

**Figure 7.**
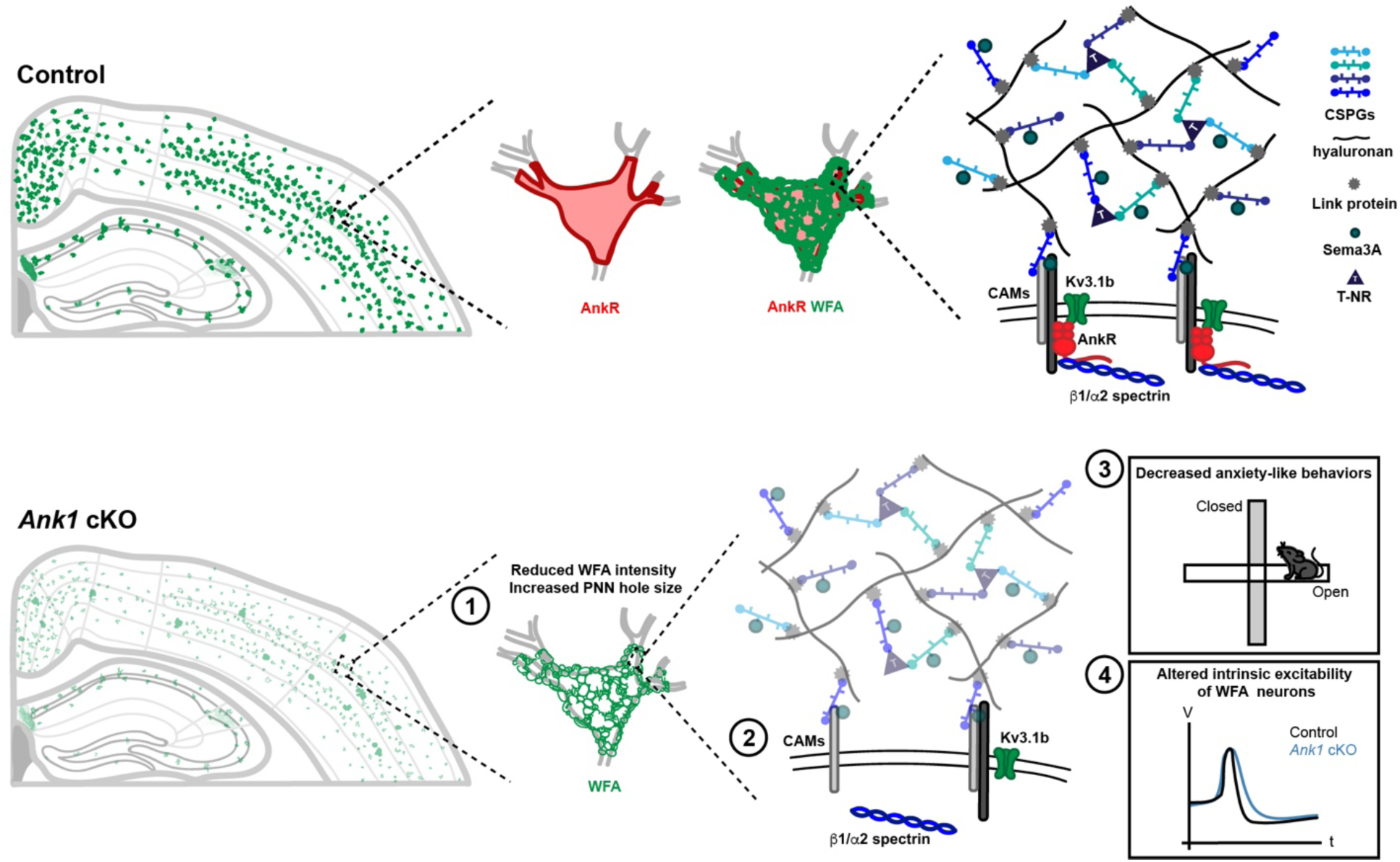
AnkR is highly expressed in the somatodendritic domain of Pv^+^ interneurons which are surrounded by PNNs, a specialized ECM structure. AnkR is a scaffolding protein that binds to and stabilizes PNN-associated CAMs (including NrCAM and PlexinA4) and ion channels (including Kv3.1b) by linking them to the *β*1-*α*2 spectrin-based cytoskeleton. Loss of AnkR results in **1)** altered PNN morphology including reduced WFA intensity and decreased compactness of the nets; **2)** molecular changes including reduced *β*1 spectrin, PNN-associated NrCAM, and Kv3.1b; **3)** behavioral changes including decreased anxiety-like behaviors in the open field and elevated plus maze; and **4)** electrophysiological changes including decreased AP latency and threshold, broader APs with shallower and delayed AHP, and decreased firing rate during current injection.

### AnkR maintains PNNs

GABAergic interneurons are surrounded by PNNs, and our proteomics experiments revealed that AnkR indirectly interacts with PNN components (**Fig. 7**). PNNs are proposed to have many functions including regulation of synaptic plasticity, excitation and inhibition, ion buffering, and even protection against neurodegeneration and neurotoxicity (Cabungcal et al., 2013; Fawcett et al., 2019; Suttkus et al., 2016b). The connection between AnkR and PNNs is remarkably similar to the connection between AnkG and perinodal and AIS ECMs. These latter ECMs interact with AnkG through the CAM NF186, and loss of either AnkG or NF186 blocks their assembly (Amor et al., 2017; Hedstrom et al., 2007; Susuki et al., 2013). In contrast, the membrane receptors and mechanisms of PNN assembly and maintenance are unknown. Our proteomics experiments revealed candidates and suggest that NrCAM and PlexinA4, together with Nrp1, may participate in assembly or maintenance of PNNs. These CAMs co-immunoprecipitate with AnkR. NrCAM and Nrp1 colocalize with a subset of AnkR^+^ neurons. Furthermore, loss of AnkR dramatically reduces the number of somatodendritic NrCAM^+^/WFA^+^ neurons, suggesting that as for PNNs, AnkR is required to maintain somatodendritic NrCAM in WFA^+^ neurons. Future studies of PNNs in NrCAM and PlexinA4-deficient mice may help to determine the role of these CAMs in PNN assembly and maintenance. In addition, other interesting candidates identified in our proteomics experiments may also function as receptors. For example, we identified the adhesion G-protein coupled receptor Gpr158 and the CAM Limbic System Associated Membrane Protein (Lsamp). It will be interesting and important to determine if these membrane proteins also function with AnkR to assemble, modulate, or maintain PNNs, and we speculate that multiple receptors link AnkR to PNNs.

We observed the disruption of the compact PNN structure and a ∼50% reduction in WFA fluorescence intensity in the absence of AnkR in juvenile and adult mice. This is notable since mice lacking four PNN components (TnC, TnR, and the CSPGs Bcan and neurocan) have a significant reduction in PNN structure, area, and WFA fluorescence during development, but PNNs normalize by postnatal day 35 (Gottschling et al., 2019). Thus, loss of AnkR has more profound effects on maintenance of PNNs than even removing components of the PNNs themselves. This suggests that although AnkR is not necessary for the assembly of PNNs, the receptors responsible for PNN maintenance converge on AnkR (**Fig. 7**). Nevertheless, since PNNs still assemble in the absence of AnkR, other AnkR-independent PNN assembly mechanisms must also exist.

### AnkR, PNNs, and psychiatric disease

Mice lacking AnkR in GABAergic interneurons showed reduced anxiety in both the open field and elevated plus maze tests. Increased anxiety has been correlated with increased PNN density (Murthy et al., 2019). Conversely, the selective serotonin reuptake inhibitor (SSRI) fluoxetine reduces anxiety as well as PNN density (Ohira et al., 2013). The results of these studies are consistent with our observation that *Ank1^F/F^;Dlx5/6-Cre* mice have both reduced anxiety and a significant reduction in PNN density (**Fig. 7**). Some studies have reported large reductions in PNNs throughout the brains of patients with schizophrenia (Berretta, 2012; Mauney et al., 2013). Multiple GWAS studies implicate *ANK1* as a schizophrenia-associated gene (Aberg et al., 2013; Fromer et al., 2014; Schizophrenia Working Group of the Psychiatric Genomics, 2014), and several of the top candidates for AnkR-interacting proteins identified in our mass spectrometry analysis (e.g. *Myh11*, *Kif1a*, *Itsn1*, and *Slc25a12*) are also schizophrenia-associated genes (Fromer et al., 2014; Schizophrenia Working Group of the Psychiatric Genomics, 2014); Kv3.1b is also reduced in patients with schizophrenia (Yanagi et al., 2014). Thus, dysregulation of *ANK1*, leading to reduced AnkR, Kv3.1b, and PNN density, may be a common pathomechanism in psychiatric disease. Intriguingly, the expression of Kv3.1b in schizophrenic patients is corrected by antipsychotic drugs (Yanagi et al., 2014). It will be interesting to determine if antipsychotic drugs similarly affect AnkR protein levels or PNNs.

### AnkR recruits Kv3.1b to the neuronal membrane

Loss of AnkR from GABAergic interneurons significantly altered their intrinsic and firing properties, suggesting disrupted K^+^ channel function. However, the changes in excitability may also reflect loss of both Kv3.1b K^+^ channels and reduced PNNs, since Bcan-deficient mice have similar changes in the excitability of their GABAergic interneurons (Favuzzi et al., 2017). We found that AnkR interacts directly with Kv3.1b and is required to maintain Kv3.1b in GABAergic neurons. We localized Kv3.1b’s AnkR-binding motif to 6 residues in its C-terminus: EDCPHI. This motif is different than the previously characterized pan ankyrin-binding motifs present in all Na^+^ channels (Garrido et al., 2003) and L1CAMs (Garver et al., 1997; Tuvia et al., 1997). In contrast, there is specificity among the ankyrin-binding capacities of different K^+^ channels: AnkR interacts with Kv3.1b and is both necessary and sufficient to induce Kv3.1b clustering in the soma of GABAergic interneurons and a subset of CNS nodes of Ranvier, while AnkG binds Kv7.2/3 and is necessary and sufficient for its clustering at AIS and nodes of Ranvier. Although the Kv7.2/3 motif is highly homologous to the Na^+^ channel ankyrin-binding motif, AnkR does not cluster Kv7.2/3 (Wang et al., 2018). When Kv3.1b is present at nodes, it is associated with AnkR. A nearly identical motif is also present in Kv3.3. Like Kv3.1b, Kv3.3 is enriched in Pv^+^ interneurons in the forebrain, hippocampus, deep cerebellar nuclei and Purkinje neurons in the cerebellum (Chang et al., 2007); these same cells also express high levels of AnkR (Kordeli and Bennett, 1991). Although one previous study suggested Kv3.3 is not found at nodes (Chang et al., 2007), our results show that CNS nodes with AnkR have both Kv3.1b and Kv3.3, most likely as heterotetramers. Thus, K^+^ channel diversity among nodes of Ranvier is dictated by the specific ankyrin scaffolds present at nodes.

### AnkR and Alzheimer’s disease

A number of epigenome-wide association studies in Alzheimer’s disease patients consistently report neuropathology-associated DNA hypermethylation of *ANK1* (De Jager et al., 2014; Higham et al., 2019; Lunnon et al., 2014; Smith et al., 2019a; Smith et al., 2019b). However, the consequence of this hypermethylation for AnkR protein expression is unknown. Experiments in APP/PS1 mice also report significant reductions in Kv3.1b (Boda et al., 2012). Since Kv3.1b levels in GABAergic neurons depend on AnkR, reduced expression of AnkR in the AD brain could result in altered neuronal excitability or circuit function due to the decreased levels of Kv3.1b. Similarly, PNN density has been reported to be reduced in both human AD brains and brains from the 5xFAD mouse model of AD (Crapser et al., 2020). Although these studies suggested that activation of microglia disrupted PNNs, our experiments show that another mechanism could be reduced expression of AnkR. PNNs are thought to protect against neurodegeneration since the loss of PNNs renders neurons more susceptible to accumulation of neurofibrillary tangles comprising phosphorylated tau, and lipofuscin (Suttkus et al., 2016a; Suttkus et al., 2016b), and neurons without PNNs are more susceptible to A*β*1-42 toxicity (Miyata et al., 2007). Thus, reduced AnkR expression may decrease PNN density, thereby rendering neurons more susceptible to injury in AD.

Since mutations in AnkR cause severe hemolytic anemia, its role in the nervous system has mostly been ignored. However, our results demonstrate that AnkR is much more than just red blood cell ankyrin. Our studies reveal AnkR’s critical role in fast-spiking GABAergic interneurons, its interacting proteins, and show that AnkR is necessary for normal interneuron function.

## ACKNOWLEDGEMENTS

The work reported here was supported by the following research grants from the National institutes of Health: R01 NS044916 (MNR), R01 GM103481 (ALB), R01 MH117089 (MX), R01 NS100893 (MX), F31 NS100300 (SRS), F30 MH118804 (CML), RF1 NS118584 (MDC). This work was also supported by the Dr. Miriam and Sheldon G. Adelson Medical Research Foundation (ALB and MNR), the HMH Clinician Scientist Award (MDC), and a Houston Methodist Hospital/Baylor College of Medicine collaborative pilot grant in Alzheimer’s disease and related dementias. CML is part of the Baylor College of Medicine Medical Scientist Training Program and McNair MD/PhD Student Scholars supported by the McNair Medical Institute at the Robert and Janice McNair Foundation. MX is a Caroline DeLuca Scholar.

## AUTHOR CONTRIBUTIONS

Conceptualization, Methodology, Validation, Data Curation, Investigation, and Writing – Original Draft and Editing, S.R.S; Methodology, Data Curation, Investigation, and Writing – Original Draft and Editing, C.M.L; Investigation L.H.T., S.N., and Y.O.; Methodology, Data Curation, Writing – Original Draft and Editing, A.S.A; Data Curation, Resources, and Writing – Review and Editing, J.O-P.; Resources, Supervision, Funding Acquisition, and Writing – Review and Editing, A.L.B., Methodology, Investigation, Funding Acquisition, and Writing – Review and Editing, M.D.C; Methodology, Supervision, Funding Acquisition, and Writing – Review and Editing, M.X.; Conceptualization, Methodology, Data Curation, Funding Acquisition, Writing – Original Draft and Editing, and Project Administration, M.N.R.

## DECLARATION OF INTERESTS

The authors declare no competing financial interests

## MATERIALS AND METHODS

### Animals

AnkR conditional knockout mice were generated using cell-type-specific Cre-mediated deletion of the *Ank1* gene. The targeting construct was designed by introducing *loxP* sites flanking exons 26 and 27 of *Ank1*, upstream of the spectrin binding domain. The Cre-mediated removal of these exons will cause a frame-shift mutation resulting in a premature stop codon in exon 28. Forward primer: 5’-GGG AAA CTC CAC AGA GCC TGA CGG GTC AGT-3’, Reverse primer: 5’-GGC GTC CCT ATG TTC CAT CCT ATA GAT GAC T-3’. Generation of the target construct, ES cell electroporation, colony selection, blastocyst injection, and generation of chimeric mice were a service of the University of Rochester Medical Center Transgenic Core Facility. The strategy is similar to that successfully used to create the *Ank2* and *Ank3* floxed mice where *loxP* sites flank exons 23/24 and 22/23, respectively(Chang et al., 2014; Ho et al., 2014). *Ank1^F/F^* mice were generated and backcrossed to C57BL/6 (IMSR Cat# JAX:000664, RRID:IMSR_JAX:000664) for at least four generations before being crossed to *Nestin-Cre* transgenic mice (IMSR Cat# JAX:003771, RRID:IMSR_JAX:003771), *Dlx5/6-Cre* transgenic mice (IMSR Cat# JAX:008199, RRID:IMSR_JAX:008199). *Ank1^pale/pale^* mice were obtained from Jackson Laboratory (IMSR Cat# JAX:009157, RRID:IMSR_JAX:009157). Since germline recombination has been found to occur in these Cre lines(Luo et al., 2020), immunostaining using anti-AnkR antibodies was used as a secondary confirmation of genotype. *Ank3^F/F^* mice (IMSR Cat# JAX:029797, RRID:IMSR_JAX:029797) were crossed with *ChAT-Cre* transgenic mice (IMSR Cat# JAX:006410, RRID:IMSR_JAX:006410). Both male and female mice were used in our studies. All experiments were conducted in compliance with the National Institutes of Health Guide for the Care and Use of Laboratory Animals and were approved by the Animal Care and Use Committee at Baylor College of Medicine.

### Blood transfusion and bone marrow transplant in *Ank1^pale/pale^* animals

The *pale* mutation in *Ank1* results in severe anemia characterized by pale skin tone at birth and death within one to two weeks of age(Ho et al., 2014). We performed a blood transfusion at P1 and subsequent bone marrow transplant at P30 which enabled *Ank1^pale/pale^* animals to survive into adulthood. Animals received a 5mL/kg body weight external jugular vein blood transfusion at P1, and 5mL/kg bone marrow transplant into the tail vein at P30. Blood was collected from the male breeding mouse and washed with two-times volume of saline prior centrifugated at 700Xg for 20 min at 4°C to remove excess plasma and debris. Bone marrow was isolated from the tibia and femur of close genealogical donor mice. In brief, bones were extracted and placed in 3mL 2% fetal bovine serum (FBS) HBSS (without Ca^2+^ and Mg^2+^). Marrow was exposed by cutting with sterile scissors and a 25 gauge needle and 3mL syringe were used to expunge the marrow into a sterile 6cm dish with 2% FBS HBSS on ice. Cells were then filtered through 1 cm^2^ 100 μm nylon mesh into a 5mL FACS tube. Cells were counted and diluted to an injection concentration of 3 million cells/100 μL volume. Throughout life, all animals were closely monitored for changes in weight and behavior.

### Behavioral Tests

Mice aged to 9-14 weeks were handled for 3 days prior to the start of testing. Animals began with the open field assessment followed by the elevated plus maze to assess locomotor activity and anxiety. The open field assessment was conducted in white acrylic open-top boxes (46 x 46 x 38cm) in a room lit by indirect white light for 30 minutes. Following this, animals were given 2-3 hours rest in their home cages. The elevated plus maze assessment was then conducted on an elevated platform for 10 minutes. Activity for these tasks was recorded and analyzed using the ANY-maze Video Tracking System version 4.99v (Stoelting Co, Wood Dale, IL).

### Antibodies

The primary antibodies used here include: mouse monoclonal antibodies against AnkR (UC Davis/NIH NeuroMab Facility Cat# 75-380, RRID:AB_2491109), *β*1 spectrin (UC Davis/NIH NeuroMab Facility Cat# 73-374, RRID:AB_2315814), AnkG (UC Davis/NIH NeuroMab Facility Cat# 73-146, RRID:AB_10697718), parvalbumin (UC Davis/NIH NeuroMab Facility Cat# 73-455, RRID:AB_2629420), actin (Millipore Cat# MAB1501, RRID:AB_2223041), tenascinR (R and D Systems Cat# MAB1624, RRID:AB_2207001), aggrecan (Millipore Cat# AB1031, RRID:AB_90460), brevican (UC Davis/NIH NeuroMab Facility Cat# 75-294, RRID:AB_2315824), NrCAM (R and D Systems Cat# MAB2034, RRID:AB_2267411), Kv3.1b (UC Davis/NIH NeuroMab Facility Cat# N16B/8, RRID:AB_2750730 and Thermo Fisher Cat# MA5-27684, RRID:AB_2735238), Kv3.3 (Antibodies-Online Cat# ABIN572016, RRID:AB_10782137), Kv7.2 (James Trimmer, University of California at Davis Cat# N26A/23, RRID:AB_2750761), Flag-tag or DDDDK-tag (MBL International Cat# M185-3L, RRID:AB_11123930); rabbit polyclonal antibodies against AnkR(Ho et al., 2014) (RRID:AB_2833096), Ank1 (Thermo Fisher Scientific Cat# PA5-63372, RRID:AB_2638015), neurofilament M (Millipore Cat# AB1987, RRID:AB_91201), parvalbumin (Novus Cat# NB120-11427, RRID:AB_791498), versican (Millipore Cat# AB1032, RRID:AB_11213831), PlexinA4 (Abcam Cat# ab39350, RRID:AB_944890), and neuropilin-1 (GeneTex Cat# GTX16786, RRID:AB_422398), Kv3.1b (Alomone Labs Cat# APC-014, RRID:AB_2040166), Kv3.3 (Alomone Labs Cat# APC-102, RRID:AB_2040170), GFP (Thermo Fisher Scientific, Cat# A-11122, RRID: AB_221569); and chicken polyclonal antibody against Neurofascin (R and D Systems Cat# AF3235, RRID:AB_10890736). Wisteria Floribunda lectins used were: Fluorescein labeled (Vector Laboratories Cat# FL-1351, RRID:AB_2336875 and Bioworld Cat# 21761065-1, RRID:AB_2833087), and Texas-Red (EY Laboratories Cat# F-3101-1, RRID:AB_2315605). Secondary antibodies were purchased from Jackson ImmunoResearch Laboratories or Life Technologies (IgG specific mouse antibodies). We encourage researchers to determine the optimal antibody dilutions for themselves as varying tissue treatment can affect staining conditions.

### Immunofluorescence

Animals were transcardially saline perfused to reduce red blood cells, then brains, spinal cords and roots were dissected and fixed in 4% paraformaldehyde (1 hour for brains and 30 min for spinal cords and roots) on ice and subsequently immersed in 20% sucrose overnight at 4°C. Tissue was embedded in Tissue-Tek OCT (Sakura Finetek 4583) mounting medium, and frozen on dry ice. Brains were sectioned at 25 μm thickness, ventral roots at 14 μm thickness, and spinal cords were sectioned at 18 μm thickness using a cryostat (Thermo Fisher Scientific Cryostar NX70). Sections were placed on 1% bovine gelatin precoated coverslips (Thermo Fisher Scientific). Sections were blocked with 10% normal goat serum in 0.1M phosphate buffer (PB) with 0.3% Triton X-100 for 1 hour at room temperature. Primary antibodies diluted in the blocking buffer and incubated at room temperature overnight, then sections were washed with blocking buffer. Secondary antibodies and lectins were incubated at room temperature for 2 hours and washed with 0.1M PB.

For immunostaining of human tissues, sections were deparaffinized and rehydrated through a series of alcohols and water. Heat-based antigen retrieval was performed using 1× antigen retrieval solution at pH 9 (Agilent Technologies; Santa Clara, CA) for 1 hour (30 min at 95C, followed by 30 min on ice). Washes with fresh phosphate-buffered saline with Tween 20 (PBS-T) were then performed and PBS with 0.3% Triton X-100 (Sigma, T8787) was applied for 10 minutes. After additional washes with PBS-T, slides were blocked with 2.5% horse serum (Vector Laboratories, Burlingame, CA) with 1% Tween 20 (ThermoFisher, BP337) and 0.1% BSA in PBS (Thermo Scientific, 37525). Primary antibody along with the above blocking solution was applied for 1-2 hours at room temperature. Secondary antibodies were applied for 1 hour at room temperature, including Alexa Fluor 555 Anti-Rabbit IgG (1:200; A21429), Alexa Fluor 555 Anti-Mouse IgG (1:200; A32727), Alexa Fluor 488 anti-Mouse IgG (1:200; A11001), and Alexa Fluor 488 anti-Rabbit IgG (1:200; A11034) (Alexa Fluor products of ThermoFisher), as appropriate. For double-labeling studies, dilutions of both primary and secondary antibodies were combined in 2.5% horse serum and applied. Slides were mounted using Vectashield Antifade mounting medium with 4ʹ,6-diamidino-2-phenylindole (DAPI; Vector Laboratories). These studies were carried out with IRB approval from Houston Methodist Hospital (Pro00010377).

### Image Analysis

Immunofluorescence labeling was visualized and images were collected on an AxioImager (Carl Zeiss) fitted with an apotome for optical sectioning, and a digital camera (AxioCam; Carl Zeiss). AxioVision (Carl Zeiss) acquisition software was used for collection of images. Images were also collected using a Nikon Eclipse Ni-E microscope fitted with a motorized X-Y stage for acquisition of fields. Stitching of images was performed using NIS-Elements (Nikon). In some instances, linear contrast and brightness adjustments were performed using Adobe Photoshop, or Z-stacks, 3D reconstruction, and cell counts were performed using NIH FIJI/ImageJ. The WFA or AnkR intensity were quantified as previously described(Gottschling et al., 2019; McCloy et al., 2014). In brief, NIH FIJI/ImageJ was used to draw an outline around each cell and circularity, area, mean fluorescence, along with several adjacent background readings were measured. The corrected total cell fluorescence (CTCF) = integrated density – (area of selected cell × mean fluorescence of background readings), was calculated. These measurements were then averaged within and across multiple brain regions. No other processing of the images was performed. Nodes in ventral roots were counted manually. Nodes were determined to have AnkG, AnkR, Kv3.1b, or Kv7.2 if there was immunofluorescence at the nodes. If there was not immunofluorescence, the node was determined to be absent of AnkG, AnkR, Kv3.1b, or Kv7.2. Human immunofluorescence preparations were reviewed by a neuropathologist (MDC). Images were captured in cellSens software 1.13 (Olympus America, Inc.; Center Valley, PA) on an Olympus BX-43 Microscope using a DP71 camera, an enhanced green fluorescent protein (EGFP) FITC/Cy2 filter cube (set number 49002, Olympus; Center Valley, PA), and a CY3/tetramethylrhodamine-isothiocyanate (TRITC) filter cube (set number 49004, Olympus). To examine the intensity and specificity of antibody labeling, slides were first examined separately under DAPI, TRITC, and FITC filters, photographed, and then merged in cellSens. All figures were assembled using Adobe Illustrator.

### Immunoblotting

Saline perfused mouse brains were homogenized in homogenization buffer (0.32M sucrose, 5 mM Na_3_PO_4_, 1 mM NaF, 0.5 mM PMSF, 1 mM Na_3_VO_4_ and protease inhibitors) in a Dounce homogenizer on ice. Homogenates were then centrifuged at 700Xg for 10 min at 4°C to remove nuclei and debris, the supernatants then underwent another centrifugation at 27200Xg for 90 min at 4°C. Pellets were resuspended in homogenization buffer and protein concentrations were measured. The samples were resolved by SDS-PAGE, transferred to nitrocellulose membrane, and immunoblotted with antibodies. Quantification of immunoblots was done using NIH FIJI/ImageJ.

### Plasmids

AnkR-GFP and *β*1 spectrin-Myc constructs were previously described(Ho et al., 2014). The full-length Kv3.1b construct was a gift from Dr. James Trimmer (University of California at Davis). To generate Flag-tagged full-length or truncated Kv3.1b constructs, parts of Kv3.1b were PCR amplified from full-length Kv3.1b and then inserted into p3XFLAG-CMV-7.1 vector. DNA constructs were verified by sequencing (Genewiz).

### Immunoprecipitation

Saline perfused mouse brains were homogenized in 20mM HEPES pH 7.4, 2mM EDTA and protease inhibitors in a Dounce homogenizer. 1% (v/v) TX-100 was added to homogenates and solubilized on a shaker for 30 min at 37°C. Lysates were then centrifuged at 700Xg for 20 min at 4°C to remove nuclei and debris, the supernatants then underwent another centrifugation at 27200Xg for 60 min at 4°C. Lysates were collected and protein concentrations were measured. The lysates used for immunoprecipitation were prepared by dilution to final protein concentration at 1 mg/ml with lysis buffer (1% (v/v) Triton X-100, 20 mM Tris-HCl pH 8.0, 10 mM EDTA, 150 mM NaCl, 10 mM NaN_3_ and protease inhibitors). Antibodies were added and samples were rotated overnight at 4°C. Protein A (polyclonal antibodies, Thermo Scientific, 20333) or Protein G (monoclonal antibodies, GE Healthcare, 17-0618-01) agarose beads were washed with 1 ml of lysis buffer three times and then rotated with the lysates for 1 hour at 4 °C. The beads were then collected and washed with 1 ml of ice-cold lysis buffer seven times and subjected to immunoblotting.

For immunoprecipitation of full-length or truncated Kv3.1b, plasmids for these proteins were co-transfected with AnkR-GFP in HEK293T cells using PEI Max (Polysciences, 24765) according to the manufacturer’s instructions. The media was replaced after 16-20 hours of transfection. Cells were lysed at 48 hours of transfection in lysis buffer (50 mM Tris-HCl, pH 8.0, 150 mM NaCl, 0.1% TritonX-100, and 1 mM EDTA with protease inhibitor), and the lysates were centrifuged at 14,000 rpm for 10 min at 4°C. Anti-GFP antibody was mixed with the supernatant and incubated overnight at 4°C. Protein G Mag Sepharose (Cytiva), was first coated with 1 mg/ml of BSA in lysis buffer for 1 hour at 4°C and washed 3 times with lysis buffer, and then incubated with the mixture of cell lysate and antibody for 1 hour at 4°C. After being washed 7 times with lysis buffer, the beads were eluted with 50 µl of 1× Laemmli sample buffer at 95°C for 5 min. The samples were analyzed by immunoblot using anti-Flag antibody.

### Mass Spectrometry

Brain homogenates were prepared as described above before lipid extraction by acetone. Chilled pure acetone was added to homogenate (4:1) then vortexed to precipitate and rotated overnight at 4°C. Samples were then centrifuged at 27200Xg for 10 min at 4°C and washed twice with chilled acetone and water (4:1). The final pellet was air dried and flash frozen on dry ice. Lysates for AnkR immunoprecipitation were prepared as described above, except the final two washes of the beads were 20mM TrisHCl pH8, 2mM CaCl_2_. The beads were then collected and flash frozen on dry ice.

For digestion, pellets of lipid extracted brain homogenates were resuspended with sonication in 8M guanidinium hydrochloride plus 200 mM ammonium bicarbonate. Protein was reduced by adding 10 mM DTT and incubating at 60°C for 30 min. After that, samples were treated with 20 mM iodoacetamide at room temperature for 30 min and digested with 2% (W/W) Trypsin/LysC mix, mass spectrometry (MS) grade (Promega) for 4 hours at room temperature. Samples were then diluted using 100 mM ammonium bicarbonate so guanidinium hydrochloride concentration was 1M and incubated at 37°C overnight. After this, another 2% W/W aliquot of the digestion enzymes was added, and the digestion was allowed to continue for 4 hours at room temperature. Digested material was recovered using SepPacks C18 cartridges (Waters), eluted in 50% acetonitrile 0.1% formic acid, evaporated and resuspended in 0.1% formic acid for mass spectrometry analysis on a QExactive Plus (Thermo Scientific), connected to a NanoAcquity™ Ultra Performance UPLC system (Waters). 2 μg aliquots of the digests were injected in a 75 μm x 15 cm PepMap RSLC C18 EasySpray column (Thermo Scientific) and peptides resolved in 90 min gradients with 0.1% formic acid in water as mobile phase A and 0.1% formic acid in acetonitrile as mobile phase B. MS was operated in data-dependent mode to automatically switch between MS and MS/MS. The top 10 precursor ions with a charge state of 2+ or higher were fragmented by HCD. A dynamic exclusion window was applied which prevented the same m/z from being selected for 30s after its acquisition

For digestion of the immunoprecipitated samples, beads were resuspended in 36 μl 10 mM DTT in 100mM NH_4_HCO_3_ and incubated for 30 min at room temperature. After this, iodoacetamide was added to a final concentration of 15 mM and samples incubated for 30 additional minutes. 0.5 μg of sequencing grade trypsin (Promega) was added to each sample and incubated at 37°C overnight. Supernatants of the beads were recovered, and beads digested again using 0.5 μg trypsin in 100mM NH_4_HCO_3_ for 2 hours. Peptides from both consecutive digestions were recovered by solid phase extraction using C18 ZipTips (Millipore), eluted in 2x7 μl aliquots of 50% MeCN 0.1% formic acid, dried and resuspended in 2.5 μl 0.1% formic acid for mass spectrometry analysis. Peptides were separated using a 75 μm x 50 cm PepMap RSLC C18 EasySpray column (Thermo Scientific) using 3 hour gradients with 0.1% formic acid in water as mobile phase A and 0.1% formic acid in acetonitrile as mobile phase B, for analysis in a Orbitrap Lumos Fusion (Thermo Scientific) in positive ion mode. MS was operated in 3 s cycles in data-dependent mode to automatically switch between MS and MS/MS, with a charge state of 2+ or higher were fragmented by HCD. A dynamic exclusion window was applied which prevented the same m/z from being selected for 30s after its acquisition

In both cases, peak lists were generated using PAVA in-house software (Guan et al., 2011). Generated peak lists were searched against the *Mus musculus* subset of the UniprotKB database (UniProtKB.2013.6.17 for the full brain samples and UniProtKB.2017.11.01 for Ip samples), using Protein Prospector (Clauser et al., 1999) with the following parameters: Enzyme specificity was set as Trypsin, and up to 2 missed cleavages per peptide were allowed. Carbamidomethylation of cysteine residues was allowed as fixed modification. N-acetylation of the N-terminus of the protein, loss of protein N-terminal methionine, pyroglutamate formation from peptide N-terminal glutamines, and oxidation of methionine were allowed as variable modifications. Mass tolerance was 10 ppm in MS and 30 ppm in MS/MS. The false positive rate was estimated by searching the data using a concatenated database which contains the original UniProtKB database, as well as a version of each original entry where the sequence has been randomized. A 1% FDR was permitted at the protein and peptide level.

### Brain slice electrophysiology

All electrophysiological experiments were performed and analyzed blind to the genotypes. Mice were anesthetized by an intraperitoneal injection of a ketamine and xylazine mix (80 mg/kg and 16 mg/kg, respectively) and transcardially perfused with cold (0–4°C) slice cutting solution containing 80 mM NaCl, 2.5 mM KCl, 1.3 mM NaH_2_PO_4_, 26 mM NaHCO_3_, 4 mM MgCl_2_, 0.5 mM CaCl_2_, 20 mM D-glucose, 75 mM sucrose and 0.5 mM sodium ascorbate (315 mosmol, pH 7.4, saturated with 95% O_2_/5% CO_2_). Brains were removed and sectioned in the cutting solution with a VT1200S vibratome (Leica) to obtain 300 μm coronal slices. Slices containing primary somatosensory cortex were collected and incubated in a custom-made interface holding chamber saturated with 95% O_2_/5% CO_2_ at 34°C for 30 min and then at room temperature for 20 min to 6 hours until they were transferred to the recording chamber. Prior to moving slices to the recording chamber, 500 μl of fluorescein labeled WFA solution (200 mg/ml, in oxygenated cutting solution) was dropped on top of slices in holding chamber and incubated for 30–45 min to label perineuronal nets. After the incubation period slices were rinsed 3-4 times with oxygenated cutting solution before transferring to the recording chamber.

Recordings were performed on submerged slices in artificial cerebrospinal fluid (ACSF) containing 119 mM NaCl, 2.5 mM KCl, 1.3 mM NaH_2_PO_4_, 26 mM NaHCO_3_, 1.3 mM MgCl_2_, 2.5 mM CaCl_2_, 20 mM D-glucose and 0.5 mM sodium ascorbate (305 mosmol, pH 7.4, saturated with 95% O_2_/5% CO_2_, perfused at 3 ml/min) at 32°C. For whole-cell recordings, a K^+^-based pipette solution containing 142 mM K^+^-gluconate, 10 mM HEPES, 1 mM EGTA, 2.5 mM MgCl_2_, 4 mM ATP-Mg, 0.3 mM GTP-Na, 10 mM Na_2_-phosphoCreatine (295 mosmol, pH 7.35) or a Cs^+^- based pipette solution containing 121 mM Cs^+^-methanesulfonate, 10 mM HEPES, 10 mM EGTA, 1.5 mM MgCl_2_, 4 mM ATP-Mg, 0.3 mM GTP-Na, 10 mM Na_2_-phosphoCreatine, and 2 mM QX314-Cl (295 mosmol, pH 7.35) was used. Membrane potentials were not corrected for liquid junction potential (experimentally measured as 12.5 mV for the K^+^-based pipette solution and 9.5 mV for the Cs^+^-based pipette solution).

Neurons were visualized with video-assisted infrared differential interference contrast imaging and WFA+ neurons were identified by epifluorescence imaging under a water immersion objective (40x, 0.8 numerical aperture) on an upright SliceScope Pro 1000 microscope (Scientifica) with an infrared IR-1000 CCD camera (DAGE-MTI). Data were acquired at 10 kHz and low-pass filtered at 4 kHz with an Axon Multiclamp 700B amplifier and an Axon Digidata 1440 Data Acquisition System under the control of Clampex 10.7 (Molecular Devices).

Neuronal intrinsic excitability was examined with the K^+^-based pipette solution. The resting membrane potential was recorded in the whole-cell current clamp mode within the first minute after break-in. After balancing the bridge, the input resistance and membrane capacitance were measured by injecting a 500-ms hyperpolarizing current pulse (50–100 pA) to generate a small membrane potential hyperpolarization (2–10 mV) from the resting membrane potential. Depolarizing currents were increased in 5- or 10-pA steps to identify rheobase currents. To generate the current-firing frequency curves, the resting membrane potential of neurons was held at –75 mV and 500-ms depolarizing current pulses were increased by 50 pA steps from 0 to 1450 pA. If depolarization block occurred prior to 1450 pA, then recording was stopped.

To record miniature synaptic currents, whole-cell voltage clamp recordings of WFA+ cells were performed with the Cs^+^-based pipette solution in ACSF containing 1 μM tetrodotoxin. mEPSCs and mIPSCs were recorded for 2–3 min at the reversal potential for inhibition (–70 mV) and excitation (+10 mV), respectively.

### Electrophysiology data analysis

Data were analyzed offline by AxoGraph X (AxoGraph Scientific). The single action potential generated by the rheobase current was used to analyze action potential characteristics. Action potential threshold was defined as the voltage at which the first derivative of voltage over time exceeded 20 V/s. The action potential latency was determined as the time between the onset of current injection and action potential threshold. Action potential amplitude was determined as the voltage difference between the action potential threshold and peak. Action potential half-width was measured as the duration of the action potential at the voltage halfway between the action potential threshold and peak. Afterhyperpolarization (AHP) amplitude was determined as the minimum voltage following the action potential peak subtracted from the action potential threshold. AHP time was determined as the time between action potential threshold and the negative peak of AHP.

To analyze spike trains, action potentials were detected using Axograph X event detection with a fixed amplitude template defined by the shape of the action potential. The current-firing frequency curves were generated by measuring the frequency of action potentials for each current injection. Spike frequency adaptation and amplitude adaptation were determined from the spike train evoked by the currents that are 2 times of the action potential threshold currents. Spike frequency adaptation was measured by determining the percent decrease between the inverse of the average of the first five inter-spike intervals and the inverse of the average of the last five inter-spike intervals. Amplitude adaptation was determined by baselining the trace at the action potential threshold for the first action potential and then calculating the percent decrease between the amplitude of the first action potential and the last one. The firing frequency within the first 100 ms of current injection was also determined. Depolarization block current was determined as the minimal current that caused the cell to reach depolarization block during the 500 ms current pulse. If depolarization block was not reached by 1450 pA current injection, 1450 pA was recorded as the value.

To detect miniature synaptic events, data were digitally low-pass filtered at 2 kHz offline and events were detected by a scaled-template algorithm (AxoGraph X). The parameters of the template for mEPSCs are: length, 5 ms; baseline, 1.5 ms; amplitude, –2 pA; rise time, 0.2 ms; and decay time, 1 ms with a detection threshold of -3.25. The parameters of the template for mIPSCs are: length, 10 ms; baseline, 3 ms; amplitude, 2 pA; rise time, 0.27 ms; and decay time, 3.7 ms with a detection threshold of 3. The integrated charge per unit time for mEPSC or mIPSC was determined by multiplying the frequency of mEPSC or mIPSC by the average charge of mEPSCs or mIPSCs, respectively.

### Statistical Analyses

No statistical methods were used to pre-determine sample sizes, but our sample sizes are similar to those previously reported(Chiang et al., 2018; Ho et al., 2014). Sets of age-matched conditional knockout mice and their controls were randomly collected from the same litter or from two litters that had close dates of birth. Data were collected and processed randomly and were analyzed using Microsoft Excel and GraphPad Prism. Except for perineuronal net quantifications, researchers were not ‘blinded’ to the conditions of the experiments for data collection and analysis. Unless otherwise stated, unpaired, two-tailed Student’s *t*-test was used for statistical analysis. Data distributions were assumed to be normal, but were not formally tested. All error bars are SEM unless otherwise indicated. Significance values indicated in figures are for post hoc comparisons.

---

**Supplemental Figure 1.**
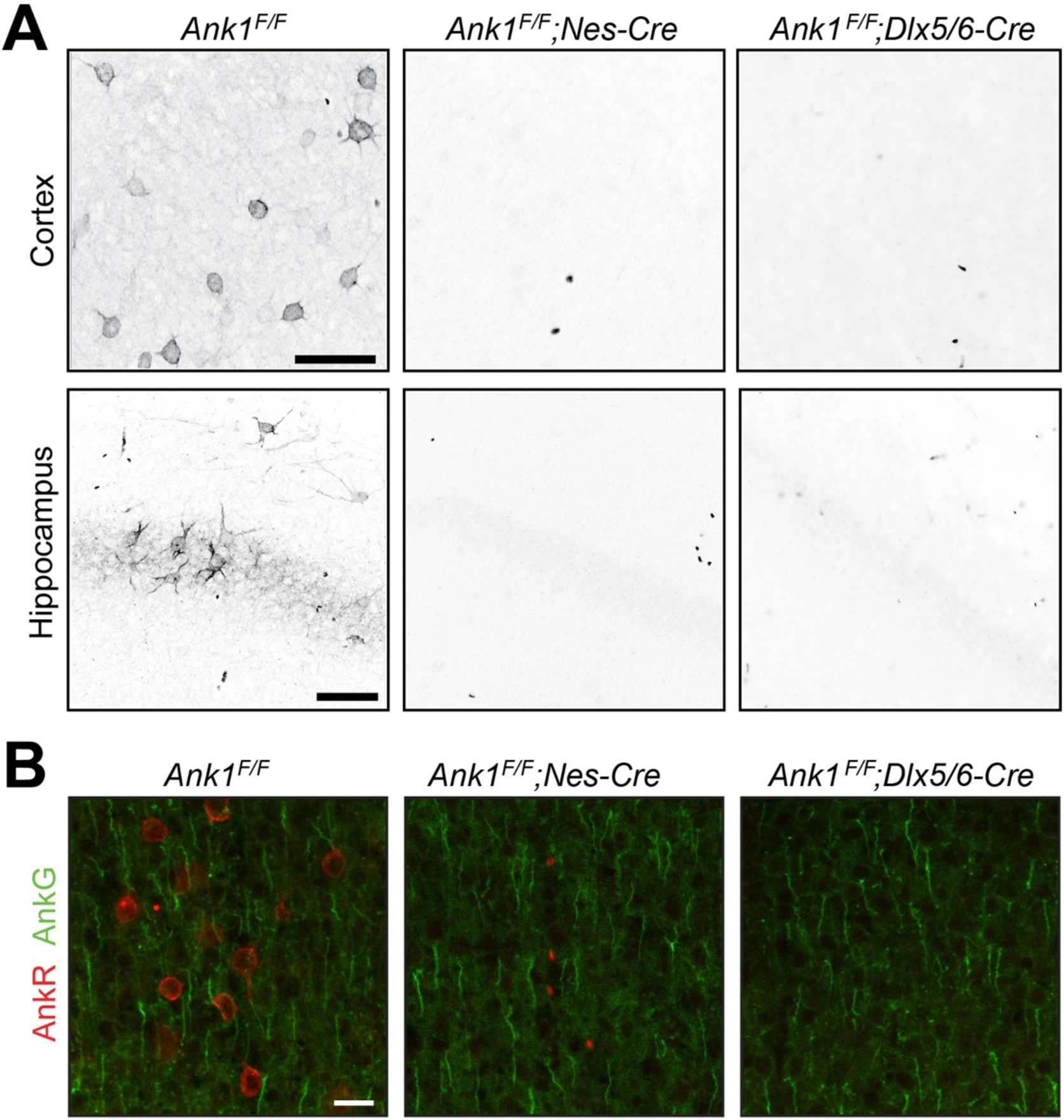
AnkR immunostaining in *Ank1* conditional knockout mice. (A) Immunostaining of cortex (top) and hippocampus (bottom) in mice with the indicated genotypes. Scalebars, 50 µm. (B) Immunostaining of cortex for AnkR (red) and AnkG (green) in the indicated genotypes. Scalebars, 20 µm.

**Supplemental Figure 2.**
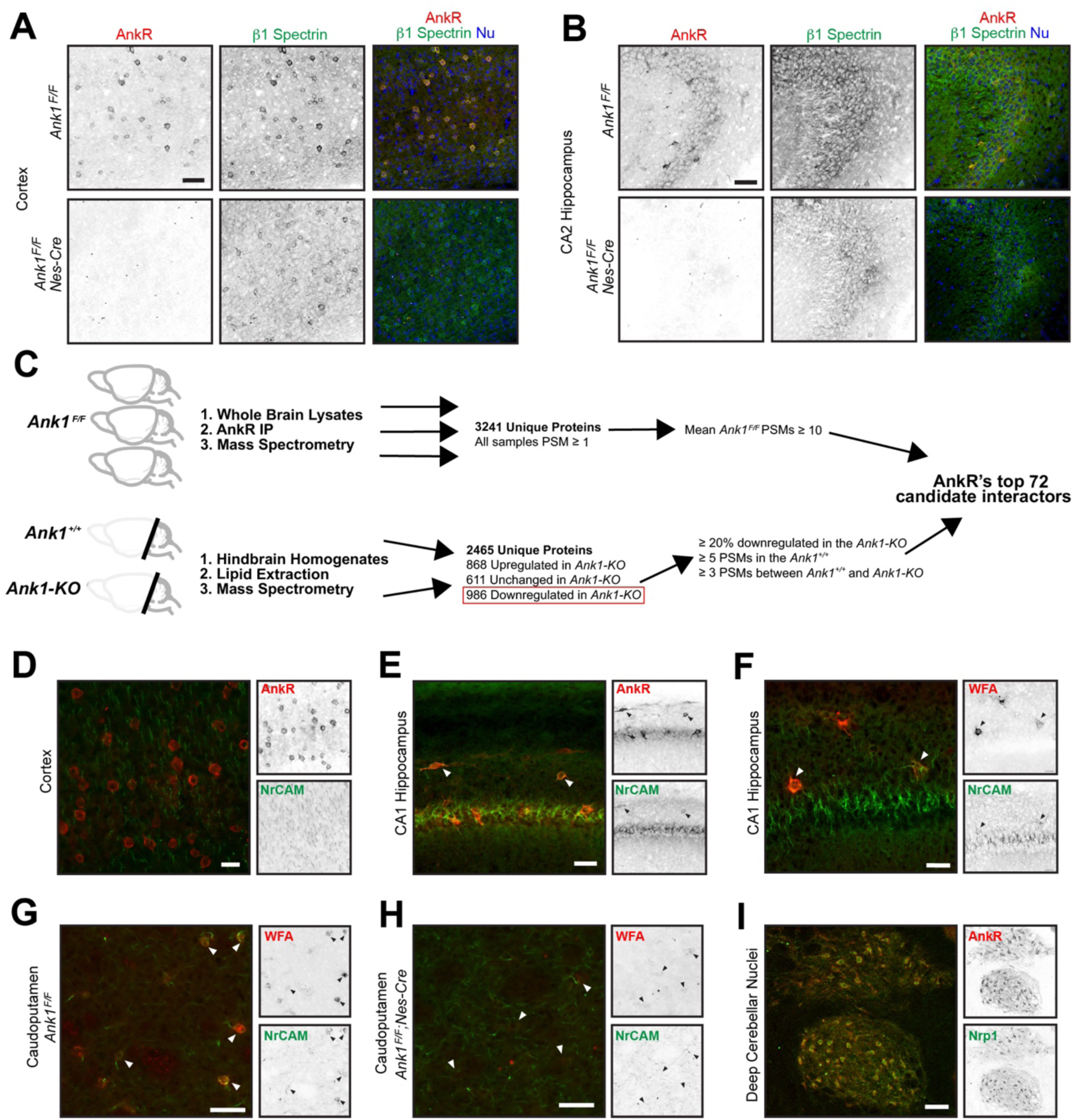
AnkR interacting proteins. (**A, B**) Immunostaining of cortex (**A**) and hippocampus (**B**) in control and AnkR-deficient mouse brain using antibodies against AnkR (red), *β*1 spectrin (green), and the nuclear marker Hoechst. Scale bars, 50 µm. (**C**) Proteomics strategy to identify AnkR-interacting proteins. PSM, peptide spectral match. (**D, E**) Immunostaining of cortex (**D**) and hippocampus (**E**) using antibodies against AnkR (red) and NrCAM (green). Arrowheads in (**E**) indicate AnkR^+^/NrCAM^+^ neurons. Scalebars, 20 µm (**D**), 50 μm (**E**). (**F**) Immunofluorescence of hippocampus using fluorescent WFA (red) and antibodies against NrCAM (green). Arrowheads indicate WFA^+^ neurons. Scalebar, 25 µm. (**G, H**) Immunostaining of caudoputamen in control and AnkR-deficient mouse brain using fluorescent WFA (red) and antibodies against NrCAM (green). Arrowheads indicate WFA^+^ neurons. Scalebars, 50 µm. (**I**) Staining of deep cerebellar nuclei using antibodies against Nrp1 (green) and AnkR (red). Scalebar, 100 µm.

**Supplemental Figure 3.**
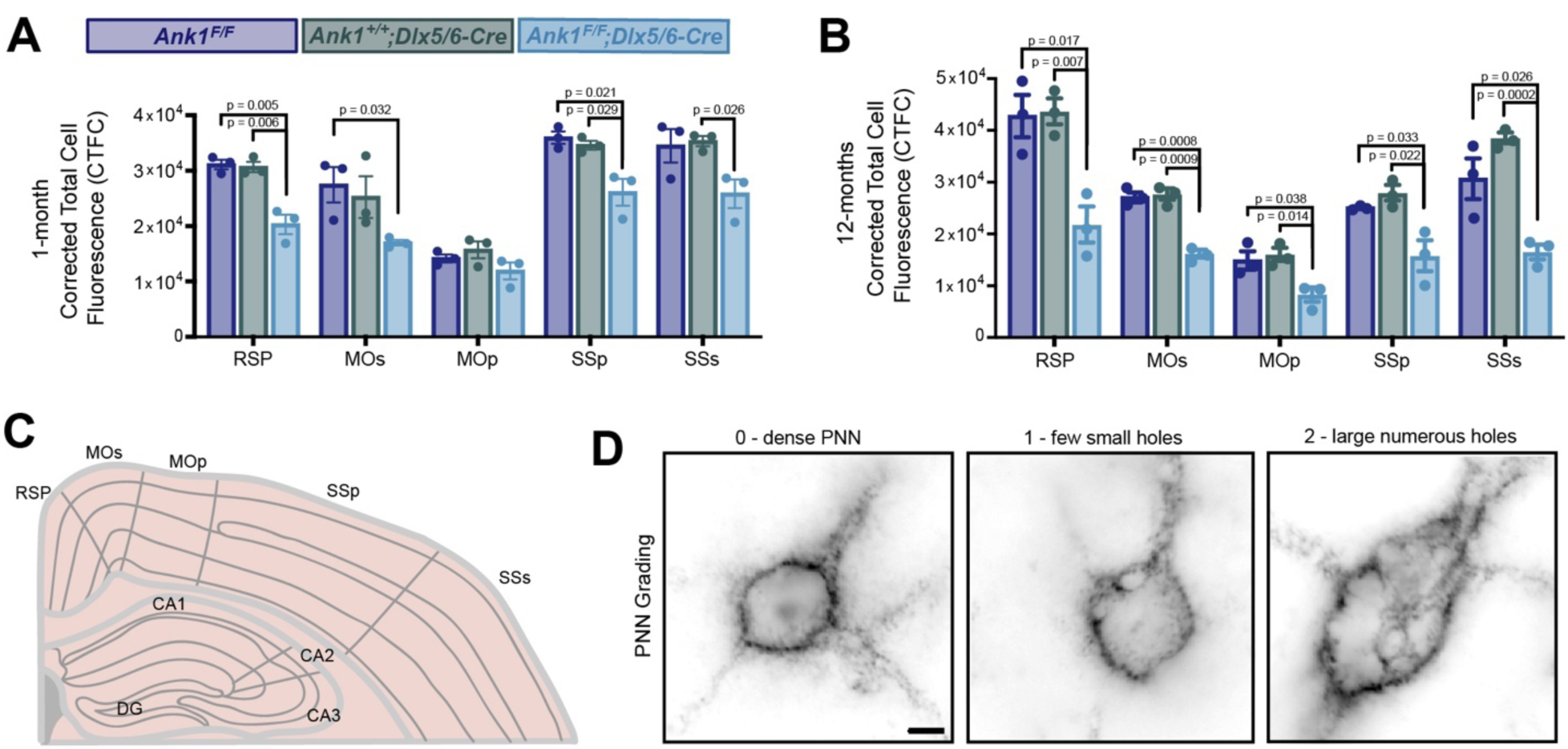
WFA labeling is reduced across cortical regions. (**A, B**) Corrected Total Cell Fluorescence (CTCF) across cortical regions at one- and 12-months of age. RSP, retrosplenial cortex; MOp, primary motor cortex; MOs, secondary motor cortex; SSp, primary somatosensory cortex; SSs, secondary somatosensory cortex. Error bars indicate mean ± SEM. N=3/group. (C) Schematic of cortical regions where CTCF was measured. (D) WFA labeling of PNNs showing grading scale used for analysis in Figures 3d and 3i. Scalebar, 5 µm.

**Supplemental Figure 4.**
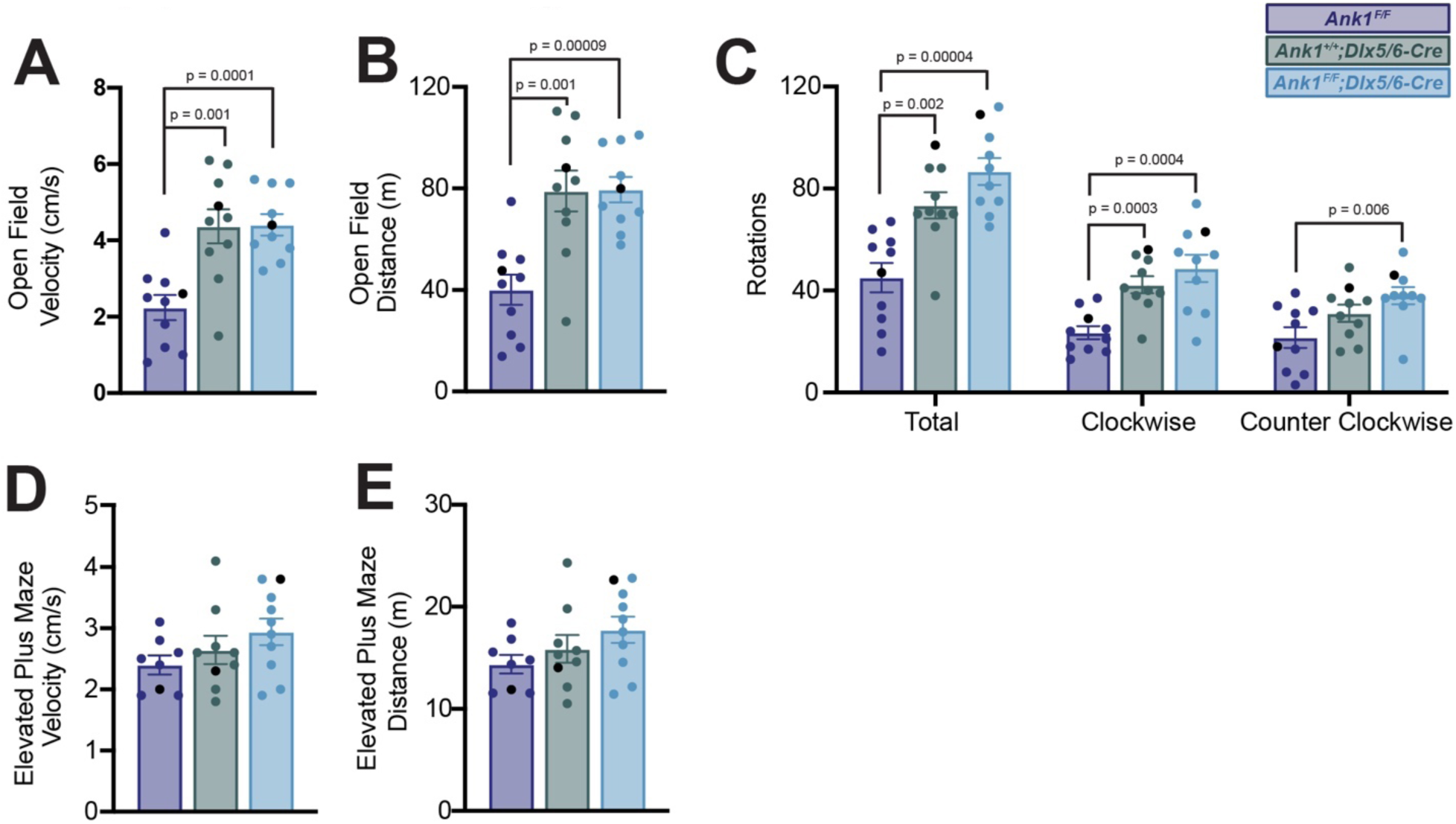
Quantification of open field and elevated plus maze assays. (**A-C**) Measurements of velocity (**A**), distance traveled (**B**), and rotations (**C**) in the open field test. (**D, E**) Measurements of velocity (**D**) and distance (**E**) in the elevated plus maze assay. In all panels error bars indicate mean ± SEM. Black circles indicate the animals corresponding to the representative traces in **Figure 4**.

**Supplemental Figure 5.**
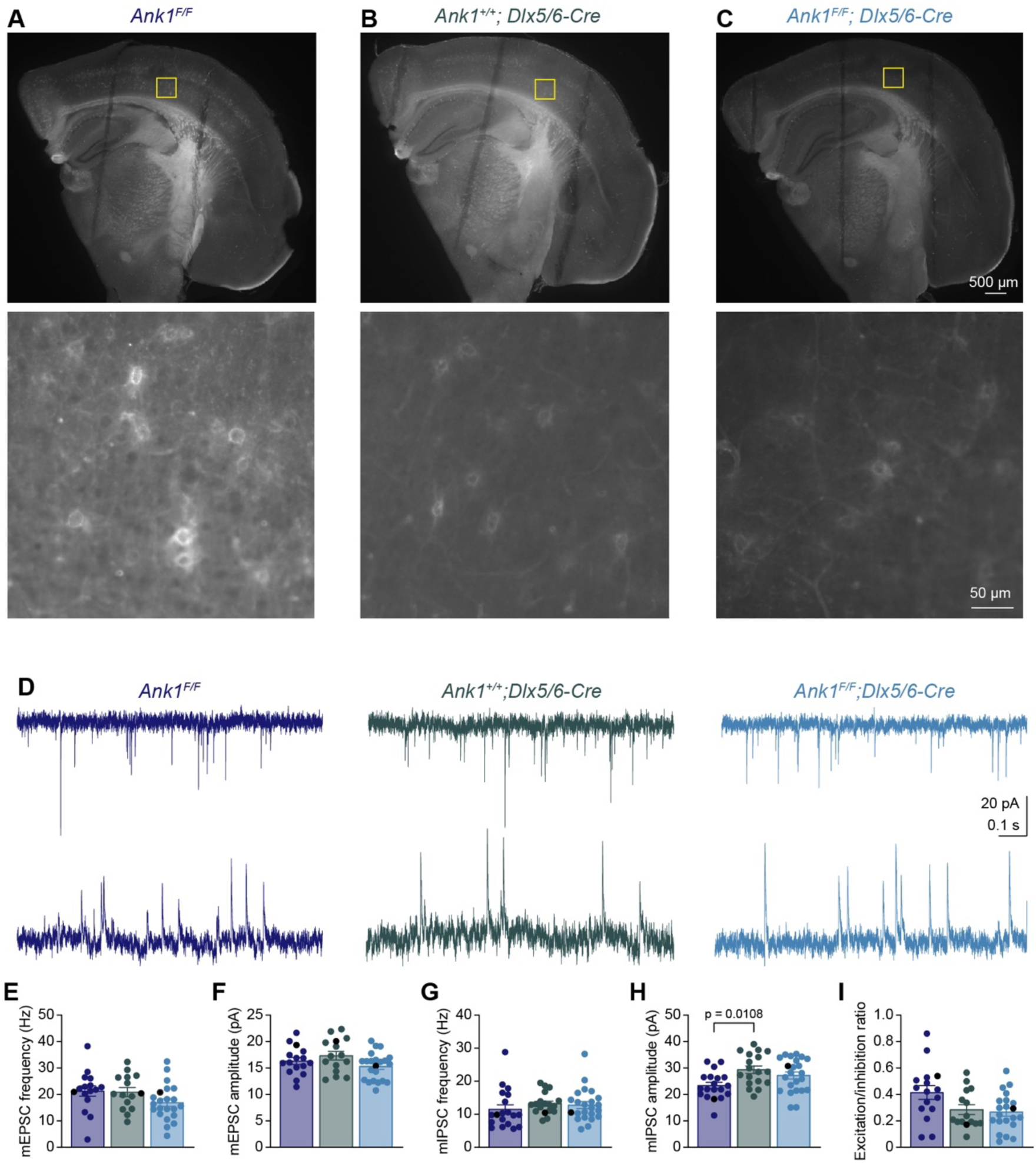
Flourescein-WFA labeling of perineuronal nets in live slices for electrophysiology. (**A**) Top: low magnification image of a coronal slice from the left hemisphere of an *Ank1^f/f^* mouse after recording. Lines on the brain slice are the indentations resulted from the mesh used to hold the slice in place during recording. Bottom: high magnification image of the somatosensory cortex layer 5 within the boxed region in the low magnification image where recording was performed. (**B, C**) Same as in (**A**) except for *Ank1^+/+^;Dlx5/6-Cre* (**B**) and *Ank1^F/F^;Dlx5/6-Cre* (**C**). (**D**) Representative traces of miniature excitatory postsynaptic currents (mEPSCs) recorded at – 70 mV (top) and miniature inhibitory postsynaptic currents (mIPSCs) recorded at +10 mV (bottom) from *Ank1^F/F^, Ank1^+/+^;Dlx5/6-Cre*, and *Ank1^F/F^;Dlx5/6-Cre* cells. (**E, F**) Summary data of mEPSC frequency (**E**) and amplitude (**F**). (**G, H**) Summary data of mIPSC frequency (**G**) and amplitude (**H**). (**I**) Summary data of excitation/inhibition ratio that is the ratio between the integrated mEPSC charge per unit time and integrated mIPSC charge per unit time. Each circle represents one neuron and the black circles indicate the representative cells in (**D**). Error bars indicate mean ± SEM. Statistical significance was determined by one-way ANOVA or Kruskal-Wallis test with multiple comparisons.

**Supplemental Figure 6.**
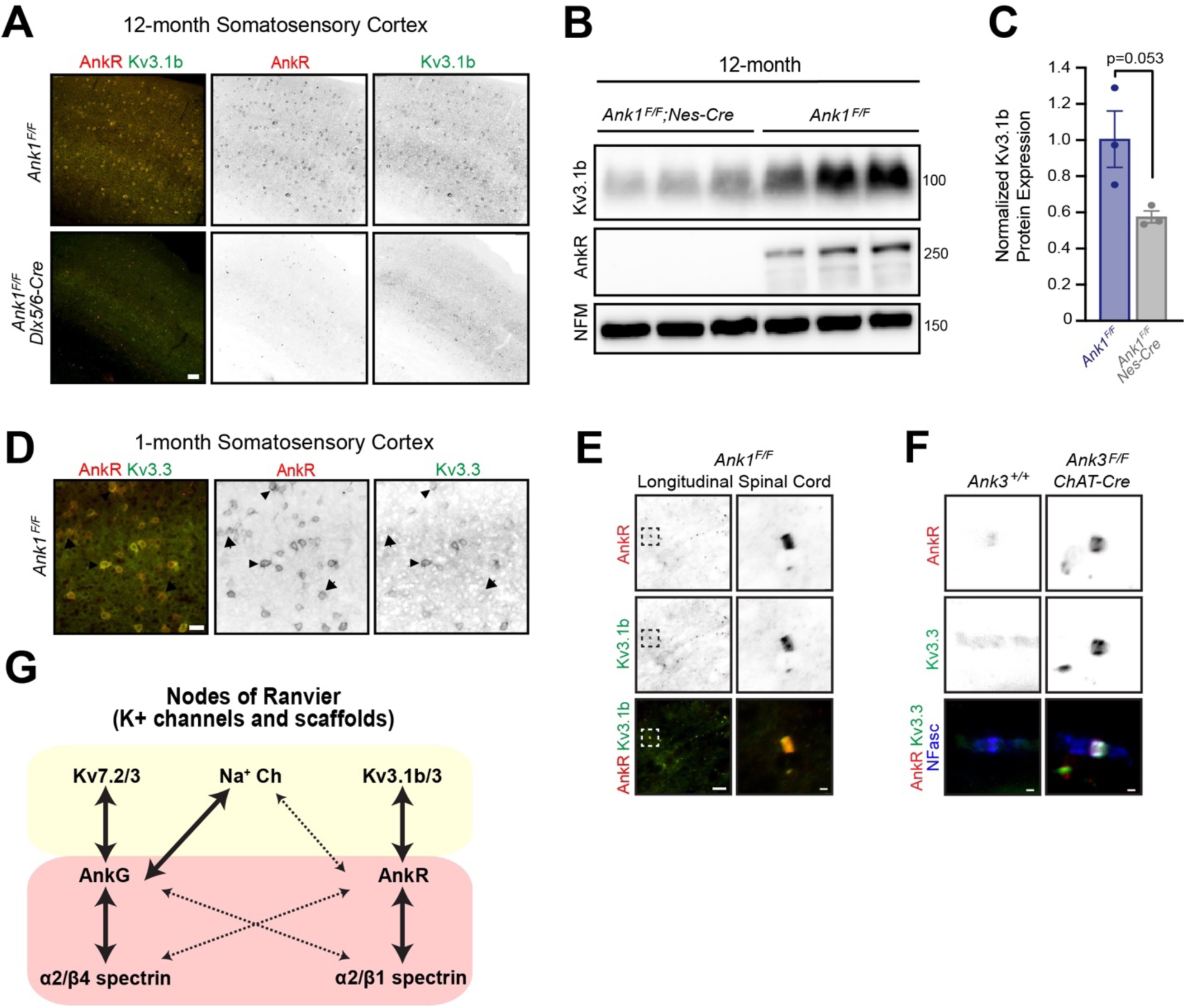
Kv3.1b membrane localization requires AnkR. (A) Immunolabeling of 12-month-old somatosensory cortex from *Ank1^F/F^* and *Ank1^F/F^;Dlx5/6-Cre* mice using antibodies against AnkR (red) and Kv3.1b (green). Scalebar, 50 μm. (B) Immunoblots of brain homogenates from three 12-month-old control and three 12-month-old AnkR-deficient brains using antibodies against Kv3.1b, AnkR, and NFM. (C) Quantification of Kv3.1b protein normalized to NFM. (D) Immunolabeling of 1-month-old somatosensory cortex from *Ank1^F/F^* mice using antibodies against AnkR (red) and Kv3.3 (green). Scalebar, 25 μm. (E) Immunostaining of *Ank1^F/F^* spinal cord using antibodies against AnkR (red) and Kv3.1b (green). Scalebars, 10 μm and 1 μm. (F) Immunostaining of ventral root nodes of Ranvier in *Ank3^+/+^ and Ank3^F/F^;ChAT-Cre* mice using antibodies against AnkR (red), Kv3.3 (green), and NFasc (blue). Scalebar, 1 μm. (G) The specific K^+^ channels and spectrins found at nodes of Ranvier is dictated by the Ankyrin scaffold to which they bind.

**Supplemental Table 1.** Intrinsic properties of WFA^+^ cells in *Ank1^F/F^*, *Ank1^+/+^;Dlx5/6-Cre*, and *Ank1^F/F^;Dlx5/6-Cre* mice. Data are from 3 *Ank1^F/F^*, 2 *Ank1^+/+^;Dlx5/6-Cre*, and 4 *Ank1^F/F^;Dlx5/6-Cre* mice, and are reported as mean ± SEM (number of cells). Bolded p values indicate significance.

|  |  |  |  | p value (One-Way ANOVA or<br>Kruskal-Wallis test) |  |  |
| --- | --- | --- | --- | --- | --- | --- |
|  | *Ank1<sup>F/F</sup>* | *Ank1<sup>+/+</sup>;*<br>*Dlx5/6-Cre* | *Ank1<sup>F/F</sup>;*<br>*Dlx5/6-Cre* | *Ank1<sup>F/F</sup>*<br>vs.<br>*Ank1<sup>+/+</sup>;*<br>*Dlx5/6-Cre* | *Ank1<sup>F/F</sup>*<br>vs.<br>*Ank1<sup>F/F</sup>;*<br>*Dlx5/6-Cre* | *Ank1<sup>+/+</sup>;*<br>*Dlx5/6-Cre*<br>vs.<br>*Ank1<sup>F/F</sup>;*<br>*Dlx5/6-Cre* |
| Resting membrane potential (mV) | -66.69 ±<br>1.373 (19) | -70.42 ±<br>0.934 (17) | -68.8 ±<br>1.102 (30) | 0.1176 | 0.403 | 0.6037 |
| Input resistance (MΩ) | 109 ±<br>10.56 (19) | 131.7 ±<br>15.94 (17) | 129.6 ±<br>7.544 (30) | 0.6384 | 0.1419 | >0.9999 |
| Membrane capacitance (pF) | 82.27 ±<br>4.849 (19) | 81.85 ±<br>25.74 (17) | 67.17 ±<br>4.068 (30) | 0.9983 | 0.0708 | 0.0956 |
| Rheobase current (pA) | 307.1 ±<br>24.56 (19) | 315.9 ±<br>26.85 (17) | 300.5 ±<br>17.75 (30) | 0.9652 | 0.9743 | 0.8768 |
| Action potential latency (ms) | 128 ±<br>39.41 (19) | 152.2 ±<br>44.77 (17) | 43.71 ±<br>18.01 (30) | >0.9999 | 0.0973 | **0.0246** |
| Action potential threshold (mV) | -33.51 ±<br>1.107 (19) | -33.09 ±<br>1.416 (17) | -37.62 ±<br>1.087 (30) | 0.9766 | **0.0165** | **0.0116** |
| Action potential amplitude (mV) | 50.88 ±<br>2.419 (19) | 53.12 ±<br>2.858 (17) | 50.41 ±<br>1.965 (30) | 0.8152 | 0.9881 | 0.6962 |
| Action potential half-width (ms) | 0.3765 ±<br>0.015 (19) | 0.3486 ±<br>0.014 (17) | 0.5316 ±<br>0.001 (30) | 0.3124 | **<0.0001** | **<0.0001** |
| Afterhyperpolarization amplitude (mV) | 26.07 ±<br>0.773 (19) | 26.9 ±<br>0.923 (17) | 19.19 ±<br>0.747 (30) | >0.9999 | **<0.0001** | **<0.0001** |
| Afterhyperpolarization time (ms) | 1.442 ±<br>0.119 (19) | 1.318 ±<br>0.087 (17) | 1.913 ±<br>0.088 (30) | 0.9943 | **0.0002** | **<0.0001** |
| Spike frequency adaptation | 0.897 ±<br>0.025 (13) | 0.9326 ±<br>0.017 (13) | 0.8912 ±<br>0.022 (24) | 0.9982 | >0.9999 | 0.721 |
| Spike amplitude adaptation | 0.8243 ±<br>0.012 (13) | 0.8446 ±<br>0.075 (13) | 0.6731 ±<br>0.027 (24) | >0.9999 | **0.0013** | **0.0001** |
| Firing frequency (Hz) | 221.5 ±<br>9.257 (13) | 329.2 ±<br>11.06 (13) | 204.6 ±<br>5.516 (24) | 0.5423 | 0.6966 | **0.0196** |
| Depolarization block current (pA) | 1438 ±<br>12.50 (8) | 1335 ±<br>59.48 (13) | 1006 ±<br>77.44 (16) | >0.9999 | **0.0031** | **0.0048** |

**Supplemental Movie 1.**
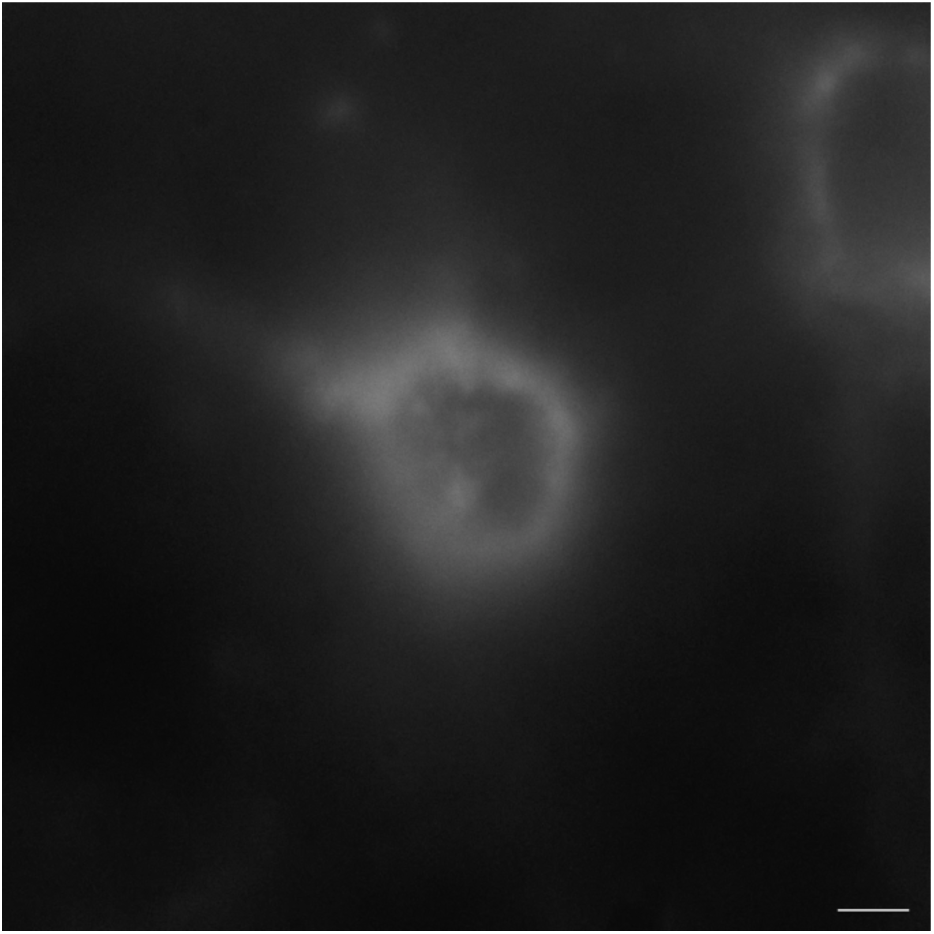
WFA in 12-month *Ank1^F/F^*. Scalebar, 5 µm.

**Supplemental Movie 2.**
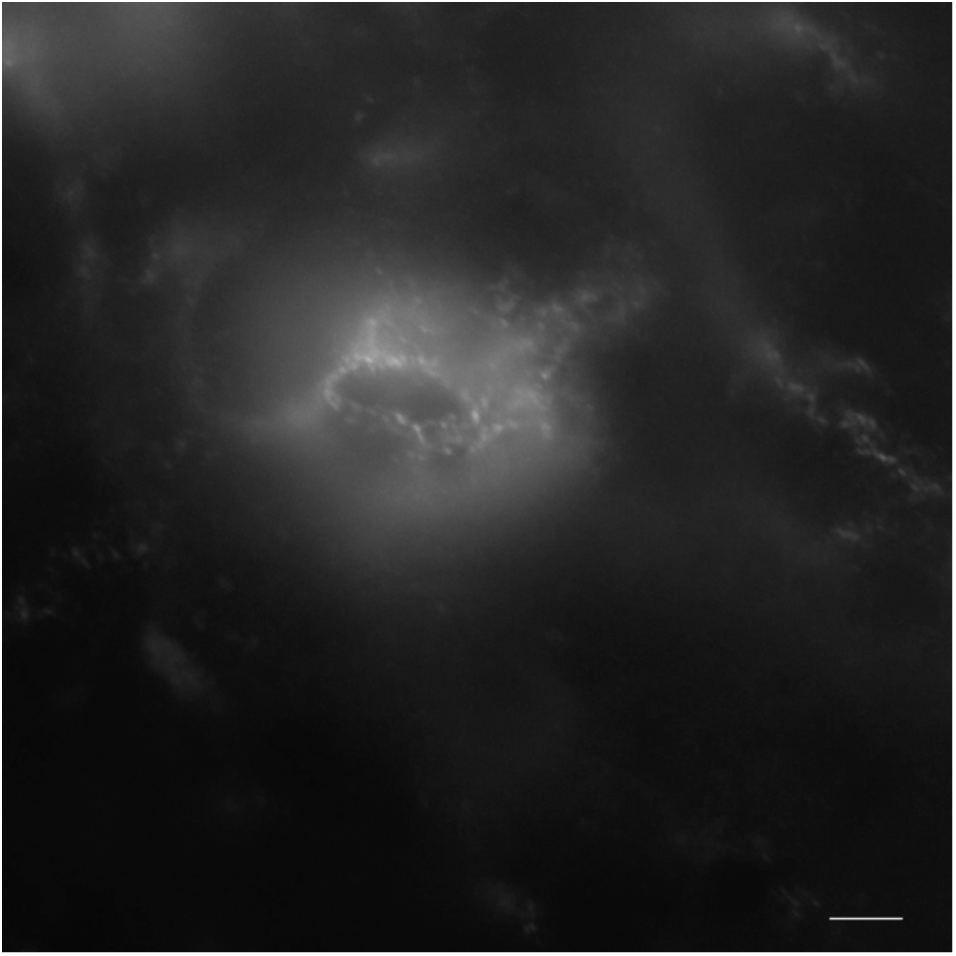
WFA in 12-month *Ank1^F/F^;Dlx5/6-Cre*. Scalebar, 5 µm.

